# Peptidoglycan turnover promotes active transport of protein through the bacterial cell wall

**DOI:** 10.1101/2025.09.12.675941

**Authors:** Zarina Akbary, Kunal Samantaray, Dan Shafir, Kaya Jain, Glen M. Hocky, Sven van Teeffelen, Enrique R. Rojas

## Abstract

The bacterial cell wall is a critical load-bearing structure, but is not thought to be an important permeability barrier since proteins freely diffuse through isolated cell wall sacculi and bacteria secrete proteins without the aid of any known channels or transporters in the wall. Using new genetically encoded probes to measure the permeability of the cell *in situ* at the single-cell level, we discovered that the size threshold determining whether proteins can pass through the *Bacillus subtilis* sacculus is smaller than was previously thought. We found that transport of small proteins (<10 kDa) through the sacculus was consistent with passive diffusion through discrete pores, while larger proteins (>15 kDa) required the generation of larger pores by inducing peptidoglycan hydrolysis unbalanced by synthesis. These data are consistent with physics-based models of diffusion through a random percolation network of finite thickness. Conversely, the ability of the innate immune factor phospholipase (15.2 kDa) to kill *B. subtilis* was inhibited by membrane de-polarization. The protective effect of de-polarization was dependent on latent peptidoglycan synthesis (decoupled from cell growth) by PBP1 – highlighting a new role for this enzyme – and on reduced peptidoglycan hydrolysis. These results demonstrate that the rapid peptidoglycan turnover that drives cell growth also promotes movement of phospholipase across the cell wall, identifying a quintessentially bacterial mechanism of active transport.

**Significance Statement:** Gram-positive bacteria, which include many serious pathogens like *Staphylococcus aureus*, *Listeria monocytogenes*, and *Clostridium difficile*, are defined by their thick peptidoglycan cell wall. Here, we demonstrate that this structure is a critical permeability barrier that blocks antibacterial proteins like those used by the human innate immune system. Furthermore, this barrier function depends on the physiological state of the cell: the wall of non-growing cells is less permeable because peptidoglycan turnover during growth actively promotes transport of specific proteins through the cell wall. This prokaryotic paradigm for molecular transport has important implications for host-pathogen interactions since pathogenic bacteria often assume both non-growing and growing states during infection.

## Introduction

The bacterial cell envelope (**Fig. 1A**) is a multi-layered structure that functions as both a permeability barrier and an exoskeleton. For Gram-negative bacteria, the plasma membrane and outer membrane are the key barriers: they are impermeable to most solutes and are therefore embedded with transporters and channels to move molecules across the envelope (1–3). In contrast, the cell wall largely determines the envelope’s mechanical properties and confers cell shape but is freely permeable to small molecules like antibiotics. The wall can block diffusion of sufficiently large macromolecules, including proteins (4), however we have only a coarse quantitative description of cell wall permeability, restricting our understanding of the functional consequences of this property. This question is particularly relevant to Gram-positive bacteria, which lack an outer membrane but possess a relatively thick cell wall that is also densely modified with teichoic acid polymers (**Fig. S1A**). Indeed, we previously observed that while most of the cytoplasm of Gram-negative bacteria rapidly diffuses through the sacculus (the isolated cell wall) when the plasma membrane is dissolved, the cytoplasm of Gram-positive bacteria appears to remain trapped by the sacculus (**Fig. 1B**) (5).

**Figure 1.**
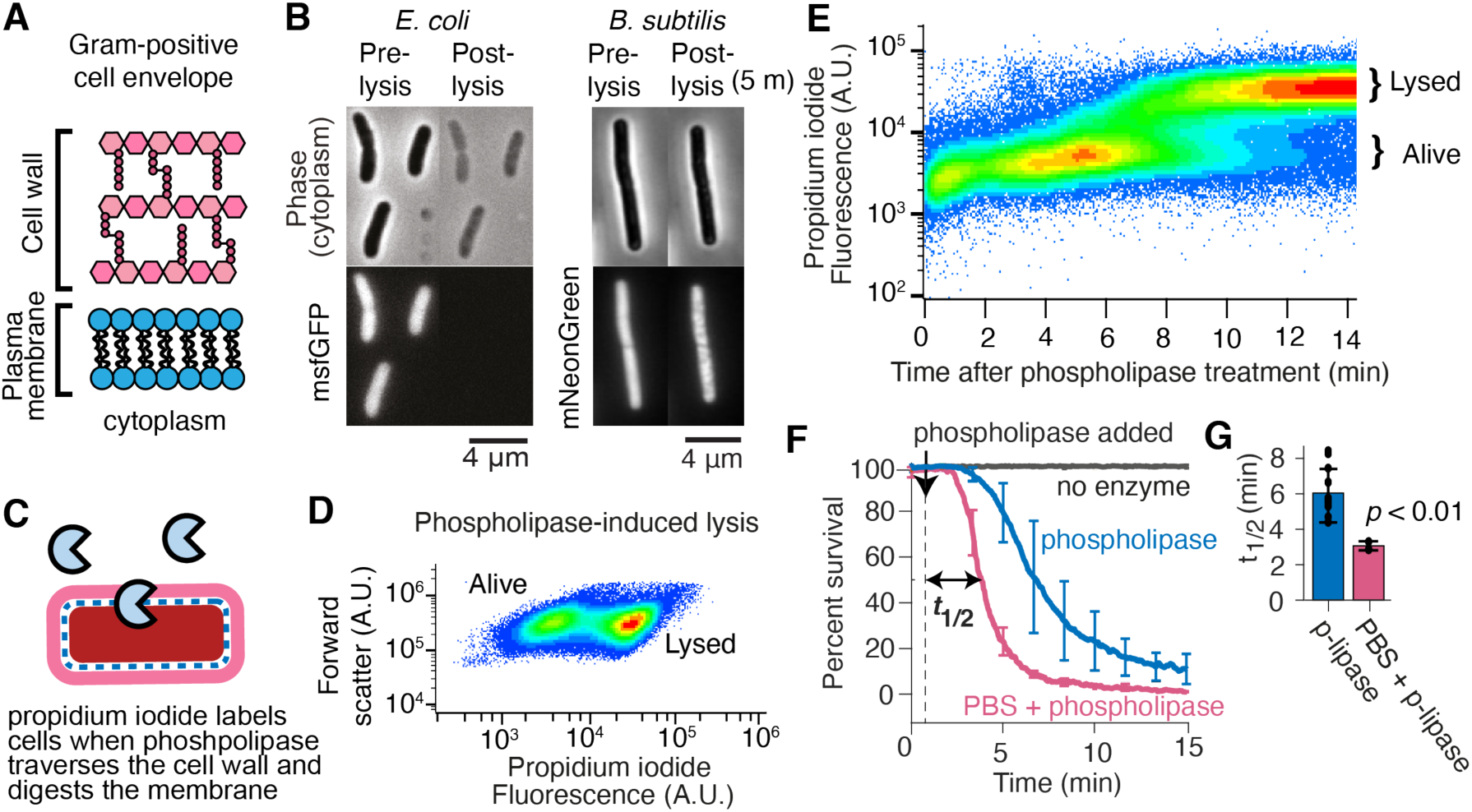
The cell wall imposes a modest permeability barrier to phospholipase. (A) Illustration of the Gram-positive cell envelope. (B) Micrographs of *E. coli* (left) and *B. subtilis* (right) expressing cytosolic GFP and mNeonGreen, respectively, before and after detergent-mediated lysis. *E. coli* data from reference (5). (*C*) Illustration of phospholipase-based assay to measure cell-wall permeability. (*D*) Cytometer forward scatter vs fluorescence of cells labeled with propidium iodide and treated with 2 μg/mL phospholipase. (*E*) Fluorescence versus time of cells labeled with propidium iodide and treated with 2 μg/mL phospholipase at *t*=0. (*F*) Mean percent survival versus time. Cells were treated with 2 μg/mL phospholipase at t = 45 seconds. Permeabilized cells were incubated in PBS for 30 minutes prior to the experiment. Error bars indicate +/- 1 s.d. across *N* = 6, 11, 3 technical replicates for no-enzyme control, phospholipase treatment, and phospholipase after permeabilization, respectively. (*G*) Mean time it takes for 50% of the cells to die, *t*_1/2_, for phospholipase treated cells and phospholipase-treated permeabilized cells. Error bars indicate +/- 1 s.d. across *N* = 6, 11 technical replicates (black dots). *p*-value is for a Student’s two-sided *t*-test.

The permeability of the cell wall will depend on its molecular structure. During the growth of rod-shaped bacteria, peptidoglycan is synthesized primarily by the multi-protein Rod complexes (**Fig. S1A**) (6–8). Drawing from a pool of membrane-anchored lipid II precursors, Rod complexes polymerize peptidoglycan oriented parallel to the circumference of the cell, on average (9–12). This anisotropic structural reinforcement explains why rod-shaped cells elongate in length without widening (13, 14). A second mode of peptidoglycan synthesis is executed by class A penicillin-binding proteins, which synthesize isotropic (non-oriented) peptidoglycan. The class A enzymes are not essential but eliminating them results in cells that are hypersensitive to antibiotics and other toxic chemicals (15). The regulation of peptidoglycan biosynthesis is not fully understood, but Rod complexes have more potential points of biochemical regulation than class A synthases, which do not function within protein complexes. For example, the scaffolding of Rod complexes by the prokaryotic actin MreB and its paralogues requires ATP hydrolysis, which directly couples this mode of peptidoglycan synthesis to global cellular metabolism, whereas the class A synthases do not hydrolyze ATP.

There are no known protein channels or transporters through the cell wall (16). Rather, the wall is thought to act as a passive “molecular sieve” with a characteristic pore size (17). As imaged with atomic force microscopy, the outer surfaces of live Gram-positive *Bacillus subtilis* and *Staphylococcus aureus* cells appear as reticulated networks of polymers with a median pore diameter of ≈40 nm, with pores as large as ≈100 nm (18). However, the inner surface of isolated cell wall sacculi appears smoother, with a median pore diameter of only ≈6 nm.

Consistent with these structural measurements, the pore size of the cell wall was estimated to be 4.12 and 4.24 nm for *Escherichia coli* and *B. subtilis*, respectively, by calculating the maximum size of dextran polymers that can diffuse into purified sacculi. This would mean that globular proteins of approximately 22 and 24 kDa are the largest that could diffuse through the sacculus. These results are surprising since they imply that the Gram-positive (*B. subtilis*) cell wall is similarly permeable to the Gram-negative (*E. coli*) wall even though the former is much thicker than the latter (19–22). Furthermore, most secreted proteins are larger than this threshold (**Fig S1B**), raising the question of how they exit the cell wall (23).

The human innate immune protein phospholipase (15.2 kDa, **Table 1**), which digests phospholipids, has been used as a molecular probe to interrogate the permeability of the cell wall in living cells (24–26) (**Fig. 1C**). Importantly, the only versions of phospholipase that killed Gram-positive bacteria (*S. aureus*, *Micrococcus luteus,* and *Listeria innocua*) are ones that had net positive charge (24). This suggests that electrostatic interactions with the cell wall—which is anionic due to its *N*-acetylmuramic acid and teichoic acid moieties—are required for phospholipase to traverse the cell wall and, furthermore, that the size threshold governing cell-wall permeability may be less for live cells (≈15 kDa) than for purified sacculi (≈25 kDa) (17).

**Table 1.** Molecular Properties of Protein Probes. R_g_: Radius of gyration.

| Protein | MW (kDa) | pI | Charge (e) | Diameter = 2R<sub>g</sub> (nm) |
| --- | --- | --- | --- | --- |
| Phospholipase | 15.2 | 8.07 | 0.638 | 4.0 |
| mNeonGreen | 26.6 | 7.06 | -0.625 | 4.4 |
| mono-ubiquitin-FIAsh | 9.5 | 5.74 | -2.167 | 3.6 |

The bactericidal activity of phospholipase also depends on the metabolic state of bacteria and is influenced by perturbations to cell wall synthesis and hydrolysis. For example, stationary-phase *S. aureus* cultures were less susceptible to phospholipase than exponentially growing cultures (25). Similarly, pre-treatment of *S. aureus* or *Enterococus faecium* with the bacteriostatic translation inhibitor erythromycin prevented phospholipase-induced lysis. These effects may depend on reduced peptidoglycan hydrolysis in non-growing cells since a mutant of *S. aureus* lacking specific cell wall hydrolases was also resistant to phospholipase. However, this mutation did not reduce the ability of the enzyme to digest phospholipids (of live cells), suggesting a synthetic lethality caused by simultaneous digestion of phospholipids and peptidoglycan rather than exclusion of phospholipase by the cell wall as a permeability barrier (25). Finally, sub-inhibitory concentrations of penicillin sensitized *S. aureus* and *E. faecium* to phospholipase (25), which is unsurprising since this antibiotic inhibits peptidoglycan synthesis.

Collectively, these observations suggest that the Gram-positive cell wall could provide a meaningful permeability barrier and that, in principle, bacteria could manipulate cell wall permeability through adaptation of peptidoglycan synthesis and hydrolysis. However, the dynamics of protein movement through the cell wall have not been measured with sufficiently high precision or temporal resolution to fully understand the mechanism(s) or functional consequences of this paradigm. Similarly, the temporal dynamics of phospholipase movement through the wall have not been decoupled from the dynamics of its enzymatic activity.

To address these questions, we constructed a panel of new fluorescent genetic probes and developed two novel assays to quantify the permeability of the Gram-positive cell wall. First, we used a cytometry-based assay to revisit the dynamics of phospholipase-induced cell lysis as a probe of wall permeability. We found that phospholipase killed exponentially growing *B. subtilis* rapidly, but that growth arrest via membrane de-polarization dramatically reduced the rate of lysis. Using a second assay to measure the efflux of mNeonGreen and fluorogenic mono-ubiquitin through the sacculus after detergent-mediated lysis, we found that depolarization-induced protection from phospholipase was not correlated with decreased permeability of the sacculus. Rather, the protective effect was dependent on continued peptidoglycan synthesis by the class A penicillin-binding protein PBP1 and on reduced peptidoglycan hydrolysis during de-polarization (27). These results demonstrate that phospholipase transport across the cell wall is an active process driven by peptidoglycan turnover.

## Results

### The cell wall imposes a weak barrier to phospholipase

In previous studies, the time for phospholipase to reduce the colony-forming units in buffer-suspensions of non-growing bacteria – roughly 30-60 minutes for *S. aureus* and *E. faecium* – was interpreted as a proxy for wall permeability (24, 26). We performed similar measurements except that we i) specifically examined phospholipase-induced lysis of *B. subtilis* growing exponentially in rich media and ii) added propidium iodide (a membrane-impermeable fluorogenic DNA dye) to the bacterial culture so that we could use cytometry to quantify cell lysis (**Fig. 1C,D**). Using this assay, we found that phospholipase killed cells rapidly, within 6.2 +/- 1.4 minutes (**Fig. 1E-G, S1C**).

To test whether the cell wall affected lysis dynamics, we subjected cells to controlled enzymatic hydrolysis of the cell wall by incubating them in phosphate buffered saline (PBS) prior to phospholipase treatment (28). PBS incubation for up to 30 minutes accelerated lysis (**Fig. 1F,G**), but longer incubation times did not have any additional effect (**Fig. S1D, E**), indicating that the basal time for phospholipase to digest phospholipids is 3.0 +/- 0.26 minutes.

We confirmed that PBS permeabilized the cell wall by developing a new single-cell microfluidics-based assay that measures the efflux of fluorescent protein probes from the sacculus after detergent-mediated dissolution of the plasma membrane (**Fig. S2A,B**). In line with our previous qualitative observations that Gram-positive sacculi retain cytoplasm after lysis (**Fig. 1B**), cytosolic mNeonGreen (**Table 1**, **Fig. 2A**) remained trapped in the sacculus for *t*lag= 27 +/- 8 minutes after lysis (**Fig. 2B-D**); after this lag, it gradually diffused out of the sacculus. However, when we pre-incubated cells with PBS to induce peptidoglycan hydrolysis, mNeonGreen efflux began immediately upon lysis at a rate that was correlated with the duration of PBS incubation (**Fig. 2E, S3A,B**). Together, our data indicate that the cell wall is a modest but not a protective barrier to phospholipase.

**Figure 2.**
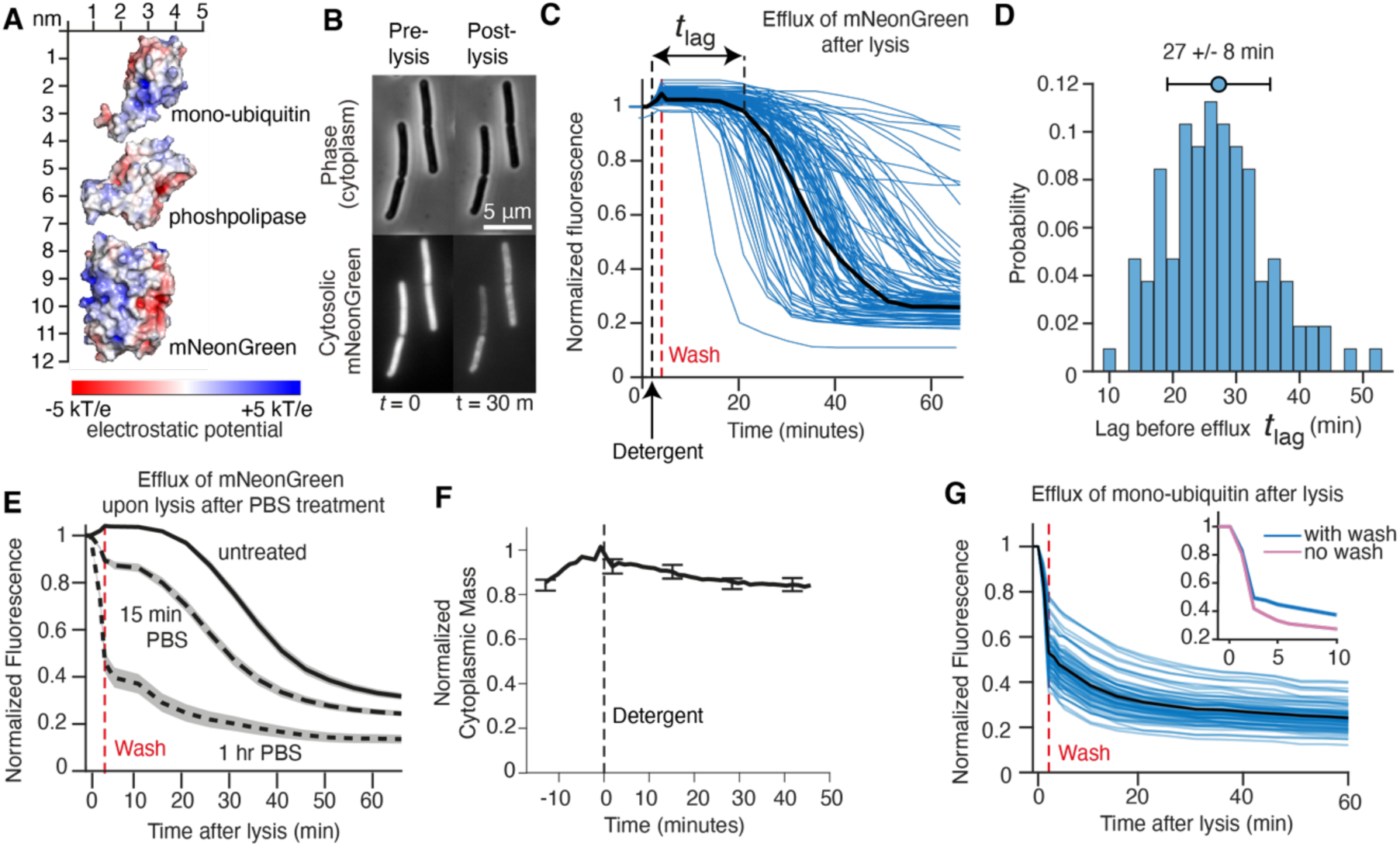
The cell wall is largely impermeable to cytoplasmic proteins. (A) Charged surface representations of mono-ubiqutin (mUbq), phospholipase (PLA2), and mNeonGreen (mNG). (B) Micrographs of *B. subtilis* before and 30 minutes after detergent-mediated lysis. *(C)* Single-cell normalized mNeonGreen fluorescence versus time from cells upon detergent treatment in microfluidic perfusion chambers (blue lines). The black line is the mean of the cell traces. The black dashed line indicates the time point at which detergent is perfused. The red dashed line indicates the time at which detergent is perfused out. *n* = 108 cells across *N* = 3 technical replicates. (*D*) The probability distribution for the lag time mNeonGreen efflux upon cell lysis (*t*_lag_). Mean (circle) and standard deviation (error bars) shown. (*E*) Mean normalized single-cell fluorescence versus time for PBS-incubated cells upon lysis. *n* = 85 and 49 cells for *N* = 1 experiment for each duration of PBS treatment. (*F*) Mean normalized single cell mass versus time for cells upon detergent treatment on an agarose pad. Error bars indicate +/- 1 s.d. from *n* = 14 cells across *N* = 1 experiment. (*G*) Normalized single-cell fluorescence versus time upon detergent treatment for cells expressing the mono-ubiquitin probe (blue lines). The black line the mean of *n* = 117 cells across *N* = 3 technical replicates. Inset: mean fluorescence from 0-10 minutes after lysis with detergent wash (blue) and without wash (pink). *n* = 170 cells across *N* = 3 technical replicates.

### The sacculus is permeable to proteins smaller than phospholipase

Our finding that the sacculus was temporarily impermeable to mNeonGreen after lysis contradicted previous studies (17, 29). To determine if mNeonGreen is unique with respect to its retention within sacculi, we performed quantitative phase microscopy to measure the total mass of the cytoplasm upon detergent-mediated lysis (30, 31). Lysis caused a rapid, small decrease in cytosolic mass, but 84% +/- 3% of the cytoplasm was retained in the sacculus after 45 minutes (**Fig. 2F**). Similarly, when we lysed cells by adding detergent to bulk liquid cultures, the optical density of the culture did not decrease for *B. subtilis* but did for *E. coli* (**Fig. S3B**). Since ribosomes and globular proteins constitute most of the cytoplasmic mass (32), our data demonstrate that the sacculus of *B. subtilis* is essentially impermeable to most of these macromolecules.

Phospholipase is slightly smaller in diameter than mNeonGreen (**Table 1**). To test if other proteins smaller than mNeonGreen can diffuse through the sacculus, we constructed a new genetic probe by fusing eukaryotic ubiquitin to the 9-amino-acid FlAsH tag (totaling 9.5 kDa), which covalently binds the membrane-permeable fluorogenic FlAsH ligand (**Fig. 2A, S4A,B**) (33). In contrast to mNeonGreen, efflux of the ubiquitin probe from the sacculus began immediately and rapidly upon cell lysis (**Fig. 2G**). Furthermore, altering its electric charge by adding a single lysine or glutamic acid residue (thereby changing the ionization equilibrium constant, pI, by one order of magnitude; **Table S1**) had no effect on its efflux (**Fig. S4C,D**). In contrast, fusions of two or three copies of ubiquitin remained trapped in the sacculus upon cell lysis, like mNeonGreen (**Fig. S4E**). Together, these data imply that mono-ubiquitin is smaller than the effective pore size of the cell wall, and that size is more important than charge for determining whether a molecule can diffuse through the cell wall.

### Protein efflux occurs through discrete pores in the sacculus

Grounded in our data, we inferred a minimal physical model explaining permeability of the sacculus after cell lysis, which was useful for interpreting further experiments. First, although the probes we used span a wide range of molecular weights, due to packing they span a very narrow range of physical sizes (**Table 1**). Therefore, our discovery that proteins smaller than mNeonGreen remain trapped in the sacculus upon lysis (**Fig. 2C,G,S4E**) implies that the maximum diameter of pores in the sacculus is less than 4.4 nm.

According to simple dimensional analysis (*Methods*), the time scale for protein efflux through the sacculus, τ, is given by

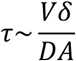

where *V* is the cell volume, *δ* is the thickness of the cell wall, *D* is the protein’s diffusion coefficient through the cell wall, and *A* is the surface area of the sacculus that the protein can diffuse through. Substituting experimentally derived parameters into **Eq. 1**, we estimated that mono-ubiquitin efflux occurs through (on the order of) 30 mono-ubiquitin-sized pores in the sacculus. Similarly, **Eq. 1** predicts that protein efflux would occur within milliseconds if the sacculus imposed no barrier to it. To test this latter prediction experimentally, we measured the dynamics of phase contrast and mNeonGreen after detergent-mediated lysis of cell wall-less protoplasts. Although we could not measure efflux at millisecond time scales, we found that both signals disappeared rapidly, within 15 seconds (**Fig. S5A**), supporting the conclusion that mono-ubiquitin efflux occurs through discrete pores in the sacculus. This experiment also demonstrates that the “slow” (≈2 min) dynamics of protein efflux are not determined by protein-protein affinity within the cytoplasm. Finally, simple theoretical arguments rule out that efflux occurs through the entire cell-wall surface but at a rate much slower than free diffusion, which predicts dynamics qualitatively different from our data (*Methods*).

It was possible that the pores we inferred were specifically engineered by the cell. For example, the number of pores per cell that we estimated is approximately equal to the number of flagella that motile cells possess, suggesting that they could be “bored” by or for flagella (34). However, mutations that resulted in constitutively motile (*Δslr*) or non-motile chaining (*ΔsigD*) cells had no effect on mono-ubiquitin efflux (**Fig. S5B,C**), arguing against this hypothesis.

In this light, we performed computational simulations of particles diffusing through finite percolating networks (35–38) to explore whether a cell wall in which peptidoglycan cross-links are randomly distributed could possess pores larger than mono-ubiquitin but none larger than mNeonGreen. In these simulations, we modeled the sacculus as a square lattice with finite thickness, δ (**Fig. 3A**). To model peptidoglycan cross-links, we randomly populated a fraction, *p,* of the discrete lattice sites inside the wall with obstacles to diffusion (*Methods*). Then, we initialized mono-ubiquitin- and mNeonGreen-sized particles at random sites within the cytoplasm, allowed them to perform Brownian diffusion, and quantified the dynamics of their efflux through the wall. Using experimentally derived parameters for these simulations (**Table S2**), we identified a critical value of *p* (*p\**) for each particle size (*Methods*): below this threshold particles rapidly diffused through pores in the sacculus, whereas above it 95% of particles remained trapped in the sacculus after 1 minute (**Fig. 3B**). Importantly, we found that there was a range *p* across which mono-ubiquitin exhibited rapid efflux through the sacculus but mNeonGreen exhibited little efflux (**Fig. 3B,C**). This analysis does not rule out a more complex mechanism of pore formation but provides proof-of-principle that a random peptidoglycan network could underlie protein efflux through the sacculus.

**Figure 3.**
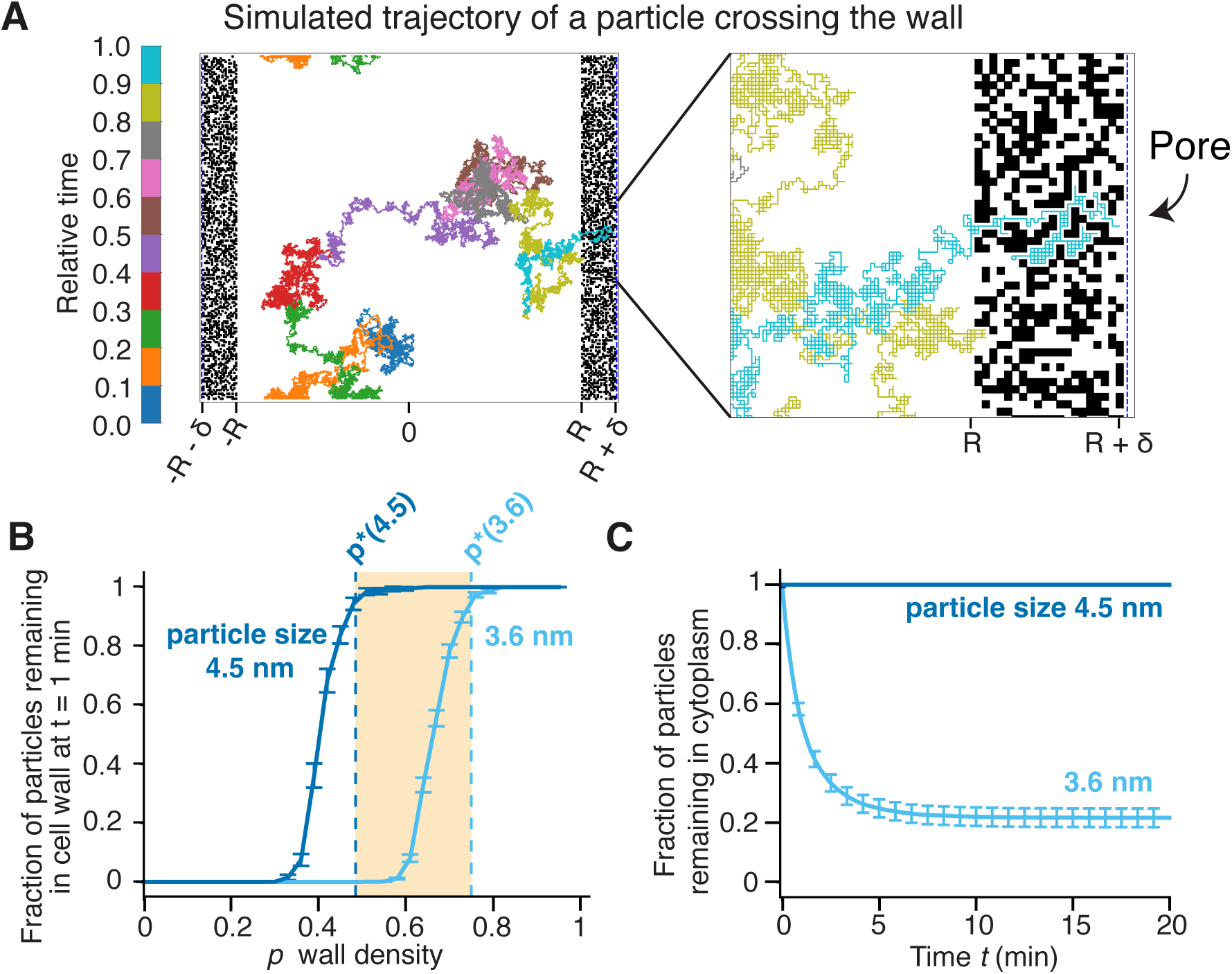
Protein efflux occurs through discrete pores. (A) illustration of simulation with an example trajectory of a particle crossing the cell wall through a pore. *R*: Cell radius. 8: Cell wall thickness. (B) The fraction of particles still inside the cell after 1 minute for 3.6 nm (light blue) and 4.5 nm (blue) particles with 8=28.8 nm. The critical values of *p* where only 5% of particles escape, indicated by the dashed lines, are *p*\* = 0.75 and *p*\* = 0.485 for a 3.6 nm and 4.5 nm-sized particles, respectively. The shaded region in between these values indicates the range of *p* where 3.6 nm particles escape and 4.5 nm particles do not. Simulation parameters are given in **Table S2**. (C) Fraction of particles that escape versus time for *p* = 0.67 and 8=28.8 nm. Error bars in (B) and (C) indicate mean +/- 1 s.d. estimated via bootstrap analysis with 200 resamples.

### Latent hydrolysis mediates efflux of mNeonGreen from the sacculus

Although mNeonGreen remained temporarily trapped in the sacculus upon cell lysis, it eventually began to exhibit efflux (**Fig. 2C**). These dynamics were not consistent with simple exponential diffusion through the cell wall, dc/dt ∝ − C. We therefore hypothesized that efflux was due to degradation of the sacculus, which could be caused by enzymatic hydrolysis of peptidoglycan.

To test this hypothesis, we measured mNeonGreen efflux after membrane dissolution in a mutant lacking the peptidoglycan hydrolases LytC and LytD, which mediate cell wall turnover and cell-cell separation after division (39, 40). These mutations resulted in a long delay in efflux (**Fig. 4A, S7A**), demonstrating that LytC and LytD largely mediate this process in wild-type cells and that these enzymes remain active within the sacculus even after cell lysis. *B. subtilis* possess 42 peptidoglycan hydrolases (41), so others are likely responsible for the slow efflux in the *ΔlytCΔlytD* mutant.

**Figure 4.**
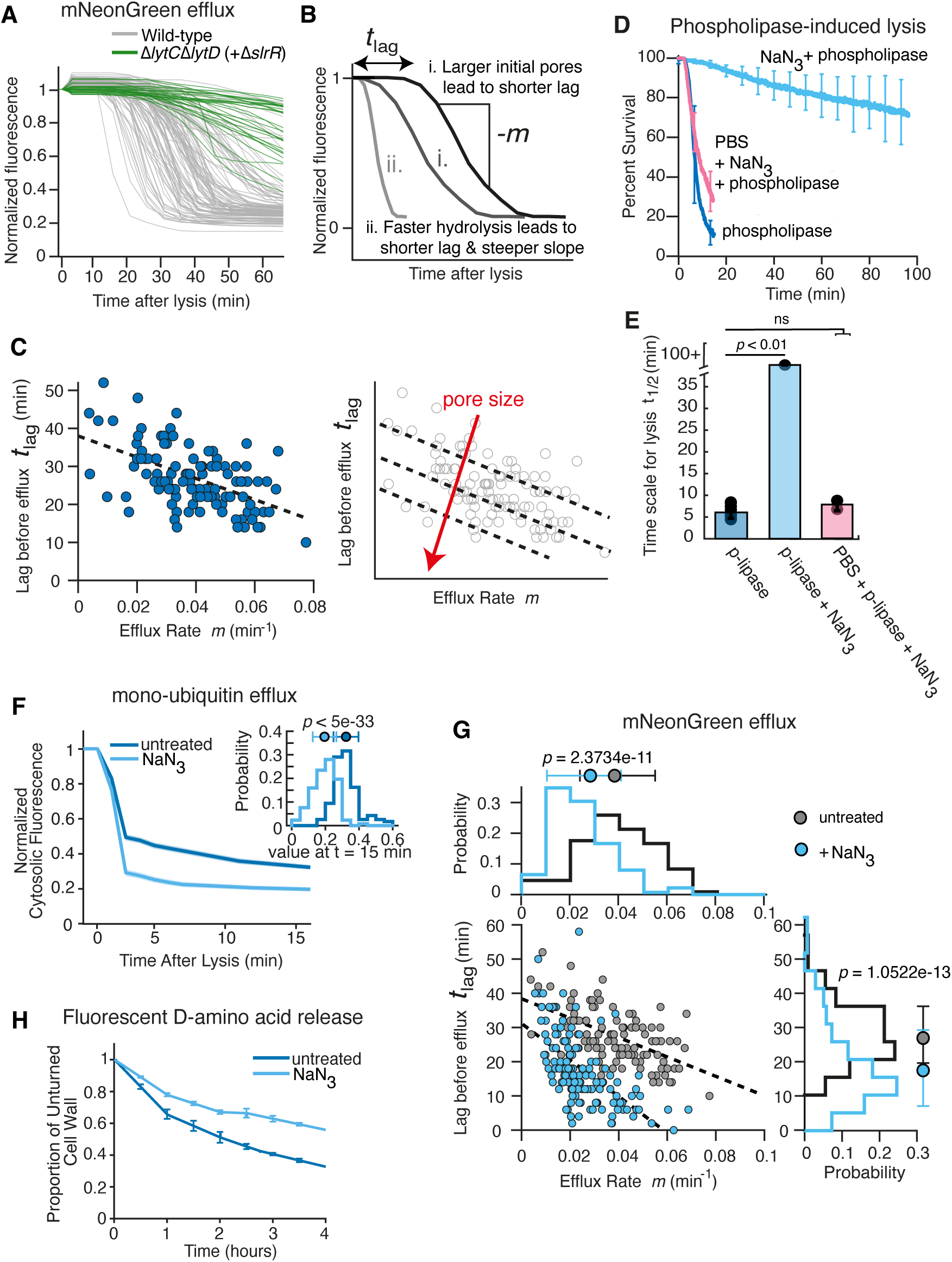
De-polarization alters cells wall permeability. (*A*) Normalized mNeonGreen fluorescence over time from wild-type (gray, from Fig. 2C) and Δ*lytC*Δ*lytD* (+Δ*slrR*) (green) cells after detergent treatment. *n* = 31 cells across *N* = 1 experiment. (B) Diagram of normalized fluorescence traces. Comparing the gray (i) trace to the black trace, the shorter lag indicates larger initial pore size but the same slope (*m*) indicates the same rate of hydrolysis. Comparing the light gray (ii) trace to the black trace, the shorter lag and steeper slope arise from a faster hydrolysis rate. (*C*) (left) Lag time before mNeonGreen efflux after lysis versus efflux rate for untreated wild-type cells shown in Fig. 2C. (right) Larger initial pore sizes at the time of lysis correspond to lower lag times for a given efflux rate. (*D*) Mean percent survival over time for wild-type cells treated with 60 mM sodium azide (light blue) one minute prior to the addition of 2 μg/mL phospholipase. Permeabilized (perm.) cells (pink) were incubated in PBS for 30 minutes prior to the experiment. *N* = 2 and 3 technical replicates for the sodium azide-treated and permeabilized sodium azide-treated traces. Wild-type + phospholipase data from Fig. 1F. (*E*) Mean time that it takes for 50% of the cells die), *t*_1/2_. The error bars indicate +/- 1 s.d. *p*-values from Student’s two-sided *t*-test, ns: *p* > 0.05. (*F*) Mean normalized single-cell fluorescence versus time after lysis for cells expressing the mono-ubiquitin probe un-treated (bue, from Fig. 2G) and treated with 60 mM sodium azide (light blue). Confidence intervals indicate +/- 1 s.e.m. *n* = 132 cells across *N* = 3 technical replicates for sodium azide-treated cells. Inset: probability distribution for the fluorescence values at t = 15 minutes for untreated (blue) and sodium-azide treated (light blue) cells. *p*-values calculated using Student’s *t*-test. (*G*) Lag before efflux versus mNeonGreen efflux rate for untreated (gray, from Fig. 2C) and sodium azide-treated (light blue) cells. Cells were treated with LB + 60 mM sodium azide for 15 minutes prior to membrane lysis. *n* = 138 cells across *N* = 3 technical replicates for sodium azide-treated cells. *p*-values calculated using Student’s *t*-test. (*H*) Fraction of fluorescent D-amino acids retained in cell wall for untreated (blue) and sodium azide-treated (60 mM, light blue) cells. Error bars indicate +/- 1 s.d. *N* = 2, 2 technical replicates.

This result revealed that the size and probably of the number of pores increases during peptidoglycan hydrolysis after cell lysis. Importantly, we discovered that 2 minutes after lysis, efflux of mono-ubiquitin and mNeonGreen after cell-wall permeabilization acutely slowed (**Fig. 2E,G**), likely corresponding to non-specific aggregation of protein in the trapped cytoplasm (**Fig. S5D**). This effect coincided with wash-out of the detergent but did not depend on it (**Fig. 2G, inset**). Protein aggregation complicated the interpretation of hydrolysis-mediated efflux of mNeonGreen since the effective size of the probe was greater after aggregation than it was before. However, we were able to use mNeonGreen as a calibrated probe before aggregation and an uncalibrated probe afterward. Accordingly, the time lag between lysis and mNeonGreen efflux is inversely dependent on the pore size prior to lysis (**Fig. 4B**). Similarly, faster hydrolysis would lead to a shorter lag and faster mNeonGreen efflux (a steeper slope associated with efflux, **Fig. 4B**). This model makes the prediction that there will be a negative correlation between the lag time, *t*lag, and the rate of efflux, *m*, across cells if there is cell-to-cell variability in the rate of peptidoglycan hydrolysis, which was validated by our experimental data (**Fig. 4C**). In other words, larger initial pores will decrease lag time for a given efflux rate (**Fig. 4C, inset**): this quantitative analysis allowed us to empirically compare pore size across bacterial mutants.

### De-polarization confers a protective effect against phospholipase

An important difference between the phospholipase-based assay for interrogating cell-wall permeability and the single-cell microfluidics-based assay is that in the former the cells are elongating and therefore phospholipase diffuses through an actively growing cell wall, while in the latter mNeonGreen and ubiquitin diffuse through the non-growing sacculus. Since both phospholipase and mNeonGreen are larger than mono-ubiquitin, and latent hydrolysis was required for efflux of mNeonGreen from the sacculus in our single-cell assay (**Fig. 4A**), we next questioned whether peptidoglycan hydrolysis during cell growth promotes transport of phospholipase through the growing cell wall (42).

To test this hypothesis, we first measured the kinetics of phospholipase-induced lysis of cells treated with sodium azide (NaN3), which de-polarizes the plasma membrane and rapidly arrests cell growth (43). Remarkably, sodium azide treatment for as little as 1 minute dramatically increased the time for phospholipase-induced lysis (**Fig. 4D, E**). To confirm that this protection was due to the permeability barrier imposed by the cell wall rather than an effect of de-polarization on the activity of phospholipase, we permeabilized the cell wall by incubating cells in PBS prior to de-polarization and the addition of phospholipase. Phospholipase killed permeabilized de-polarized cells as fast as it killed permeabilized polarized cells (**Fig. 4D, E**), demonstrating that the cell wall of de-polarized cells is impermeable to phospholipase.

We considered two non-mutually exclusive explanations for why de-polarization protected cells from phospholipase. First, de-polarization could structurally “seal” the cell wall by increasing the ratio of peptidoglycan synthesis-to-hydrolysis, thereby decreasing wall permeability. Alternatively, de-polarization, which de-energizes cells, could prevent active transport of phospholipase across the cell wall.

The first model made the prediction that de-polarization not only decreases the permeability of the living cell wall to phospholipase, but also decreases the permeability of the sacculus. To test this, we measured the effect of pre-treating cells with sodium azide on mono-ubiquitin and mNeonGreen efflux upon cell lysis. Surprisingly, de-polarizing cells for 15 minutes prior to cell lysis slightly increased the permeability of the sacculus to both probes (**Fig. 4F,G, S7B,C**): mono-ubiquitin exhibited faster and more complete efflux and, although the sacculus was still temporarily impermeable to mNeonGreen, the lag before efflux decreased. These data do not support a model in which de-polarization decreases permeability of the cell wall to phospholipase by sealing phospholipase-sized pores. Rather, since permeability of the sacculus increases during de-polarization, when phospholipase is unable to lyse cells, our data supported a model in which there are no phospholipase-sized pores either before or after de-polarization. This also implied, indirectly, that de-polarization inhibits phospholipase because its transport across the cell wall is active.

### Peptidoglycan hydrolysis and PBP1-mediated synthesis continue during de-polarization

We hypothesized that active transport of phospholipase during cell growth was directly promoted by peptidoglycan hydrolysis. The increased permeability of the sacculus to mono-ubiquitin and mNeonGreen caused by de-polarization suggested that this treatment did not completely stop hydrolysis (**Fig. 4F,G, S7B,C**). However, since de-polarization arrested cell growth, we reasoned that peptidoglycan hydrolysis was slower in de-polarized cells, and we hypothesized that slower hydrolysis led to slower transport of phospholipase across the cell wall, and therefore slower lysis. To explicitly measure how de-polarization affected hydrolysis, we covalently labeled the cell wall with fluorescent D-amino acids and used cytometry to measure the rate at which they were released as a proxy for hydrolysis (45). We found that the label was released in de-polarized cells at a rate of 0.16 h^-1^ +/- 0.01 h^-1^, which was approximately half the rate at which it was released from exponentially growing cells, 0.30 h^-1^ +/- 0.01 h^-1^ (**Fig. 4H**).

If peptidoglycan hydrolysis, unbalanced by synthesis, continues during de-polarization we would not expect this perturbation to protect cells from phospholipase. We therefore hypothesized that peptidoglycan synthesis also continues during de-polarization. Although we previously demonstrated that de-polarization causes rapid dissociation of Rod complexes (43), we reasoned that PBP1 activity might continue. Consistent with this hypothesis, elimination of PBP1 by genetic deletion of *ponA* greatly reduced the protective effect of de-polarization (**Fig. 5A,B**). De-polarization did provide modest protection of the *ΔponA* mutant against phospholipase compared to untreated *ΔponA* cells (**Fig. 5A,B**): according to our model, this would result from the reduced rate of peptidoglycan hydrolysis during de-polarization (**Fig. 4H**). Finally, simultaneously treating cells with sodium azide and vancomycin, which inhibits all peptidoglycan synthesis, resulted in precisely the same phospholipase-induced lysis dynamics that sodium azide caused in the *ΔponA* mutant (**Fig. 5C**), demonstrating i) that the activity and not just the presence of PBP1 protects cells from phospholipase during de-polarization ii) PBP1 is the main enzyme synthesizing peptidoglycan during de-polarization.

**Figure 5.**
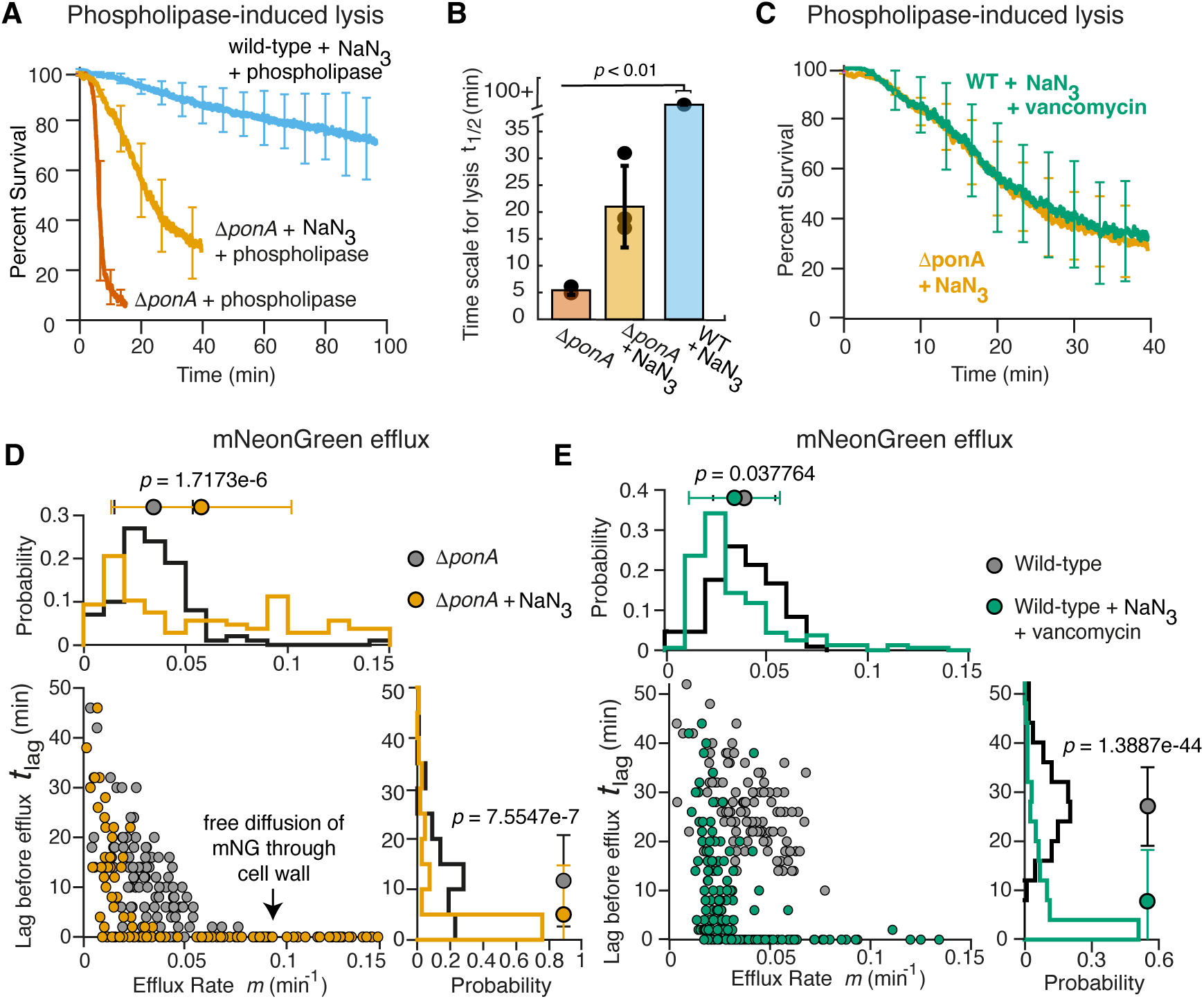
PBP1 uses latent metabolic flux to protect cells during de-polarization. (*A*) Mean percent survival over time for Δ*ponA* untreated (red) and Δ*ponA* cells treated with 60 mM sodium azide one minute prior to the addition of 2 μg/mL phospholipase (orange) and wild-type cells treated with azide (cyan, data from Fig. 1D). *N* = 2, 2 technical replicates for Δ*ponA* experiments. (*B*) Mean time it takes for 50% of the cells die, t_1/2_. The error bars indicate +/- 1 s.d. *p*-values calculated using Student’s two sided *t*-test, ns: p > 0.05. (*C*) Mean percent survival over time for wild-type cells treated with 60 mM sodium azide + 10 μg/mL vancomycin (green, *N* = 2 technical replicates) or Δ*ponA* cells treated with sodium azide (orange, data from *A*) one minute prior to the addition of 2 μg/mL phospholipase. (*D*) Time lag between lysis and efflux versus mNeonGreen efflux rate for Δ*ponA* (gray) and Δ*ponA* + sodium azide (orange) cells. Cells were treated with LB + 60 mM sodium azide for 15 minutes prior to membrane lysis. For Δ*ponA* untreated, *n* = 100 cells across *N* = 3 technical replicates. For Δ*ponA* + sodium azide, *n* = 107 cells across *N* = 3 technical replicates. (*E*) Time lag between lysis and efflux versus mNeonGreen efflux rate for untreated wild-type (gray, from Fig. 4C) and sodium azide + vancomycin-treated cells (green) cells. Cells were treated with LB + 60 mM sodium azide + 10 μg/mL vancomycin for 15 minutes prior to membrane lysis. *n* = 161 cells across *N* = 3 technical replicates for sodium azide + vancomycin-treated cells.

Based on these results, we hypothesized that de-polarization of *ΔponA* mutant cells would result in a more porous cell wall than untreated *ΔponA* cells possess. Indeed, de-polarization induced large pores in the *ΔponA* sacculus to the point where, for a fraction of cells, mNeonGreen efflux was consistent with rapid diffusion through pre-existing pores rather than latent hydrolysis-mediated efflux (**Fig. 5D, S7D**). Similarly, simultaneous de-polarization and vancomycin treatment also led to large initial mNeonGreen-sized pores in the sacculus (**Fig. 5E, S7E**).

In sum, hydrolysis and PBP1-mediated synthesis of peptidoglycan continues during de-polarization, but since both processes are slower than for growing cells, active transport of phospholipase is also slowed, leading to slower lysis dynamics (**Fig. 5F**).

### PBP1 slightly decreases the permeability of the cell wall

It has been proposed that PBP1 fills in “gaps” in the cell wall left by Rod complexes (27, 46); this motivated us to test whether we could measure these gaps as an increase in cell wall permeability. It is important to keep in mind, however, that deletion of *ponA* results in global transcriptional changes that include the upregulation of Rod complex components (8), which is likely to partially compensate for the lack of PBP1-mediated synthesis. In line with this, deletion of *ponA* had no effect on the time scale of phospholipase-induced lysis (**Fig. S8A**) or mono-ubiquitin efflux (**Fig. S8B**). Absence of PBP1 did lead to lower lag times before hydrolysis-mediated mNeonGreen efflux, indicating larger initial pores (**Fig. S8C, D**). However, these pores were still not large enough to allow free diffusion of mNeonGreen through the sacculus. These results demonstrate that deletion of *ponA* has only a modest effect on cell wall permeability and suggests that one consequence of the regulatory changes caused by the absence of PBP1 may be to ensure that permeability of the cell wall remains low.

## Discussion

Together, our analysis presents a complex new model of molecular transport through the Gram-positive cell wall. According to this model, structurally, the cell wall is impermeable to most proteins, except through relatively few pores through which small proteins (<15 kDa) can diffuse passively (**Fig. 2G**). During cell growth, however, peptidoglycan turnover (coordinated peptidoglycan synthesis and hydrolysis) can promote active transport of moderately larger proteins through the wall (**Fig. 5F**).

Atomic force microscopy of the inner surface of isolated sacculi is consistent with our structural model: the wall appears as a dense network with nanometer-scale features, punctured sparsely with large pores in the 10 nm diameter range (18). The large pores are not uniform, so it is unclear why they would only allow ubiquitin, but not mNeonGreen to pass through them. An important insight from our study that may answer this question is that as soon as growth is arrested, and even after cells are lysed with detergent, ongoing peptidoglycan hydrolysis immediately alters cell wall structure. We hypothesize that what were monodisperse ubiquitin-sized pores during cell growth become pores with larger, varying sizes during sample preparation.

It was surprising to us that the sacculus was readily permeable to mono-ubiquitin but essentially impermeable to mNeonGreen since these proteins are less than 1 nm apart in size (**Fig. 2**, **Table 1**). Our simplistic theoretical analysis based in percolation theory, however, revealed that random spatial peptidoglycan synthesis and hydrolysis could, in principle, underlie these observations (**Fig. 3**). We find it likely that feedback between pores and peptidoglycan synthesis additionally refines this pore structure. Indeed, in concurrent work, we discovered that mono-ubiquitin-sized holes are partially blocked by teichoic acids, and that eliminating teichoic acids activates PBP1 (47). This finding highlights the functional consequences of the cell wall’s role as permeability barrier. Furthermore, PBP1 activation was dependent on its terminal intrinsically disordered domain, which is approximately the molecular weight of mono-ubiquitin (**Table 1**). This mechanism would explain why the maximum pore size is mono-ubiquitin-sized if larger pores are rapidly filled by PBP1-mediated synthesis.

A major open question resulting from our study is whether there are other proteins besides phospholipase that can move actively through the cell wall: what range of protein size and charge allows this mechanism of active transport? Since most bacterial proteins are larger than ubiquitin, it is plausible that bacteria take advantage of this mechanism to promote protein secretion through the cell wall. However, from our analysis we still do not know whether the active cell wall is permeable (in either direction) to proteins larger than phospholipase (e.g. mNeonGreen). Similarly, previous studies suggest that the positive net charge of phospholipase is required for its ability to lyse cells (24), meaning that this property may be required for its active transport, for example, by promoting binding to the cell wall. However, this needs to be tested explicitly and new probes will be required to do so.

Similarly, smaller probes will be required to probe the fine-scale pore structure across most of the cell wall. Several antibiotics in the 1-2 kDa range (e.g., vancomycin, moenomycin) are rapidly effective against *B. subtilis,* and therefore the cell wall is probably freely permeable to them. This would place the size threshold governing diffusion through this phase of the cell wall in the 2-9 kDa range (roughly 1-3 nm), which is much smaller than previously predicted (17).

From an applied science standpoint, we anticipate that developing a more complete set of molecular probes to measure cell-wall permeability in a wider diversity of bacteria will enable us to functionally manipulate cell wall permeability for applications in medicine and bioengineering. For example, in the future we may be able to biochemically tailor designer protein toxins to target specific Gram-positive bacteria.

## Materials and Methods

### Bacterial growth conditions

Overnight cultures were grown by inoculating Luria Broth (LB) with frozen stock and incubated at 37 C with shaking at 180 RPM.

### Bacterial strain construction

**ER565** harboring amyE::Phyperspank-ubiquitin-TC::erm was constructed by performing a three-piece Gibson assembly reaction that contained the following PCR products and transforming into ER475. (1) a 4.7 kb fragment containing the sequence upstream of *amyE*, the erythromycin resistance cassette, and the Phyperspank promoter amplified from ER300 genomic DNA using oZA005 and oZA021 (2) a 300 bp fragment ordered from IDT, oZA006 (3) a 1.2 kb fragment containing the sequence downstream of *amyE* amplified from ER300 genomic DNA using oZA002 and oZA022.

**ER568** harboring amyE::Phyperspank-ubiquitin-ubiquitin-TC::erm was constructed by performing a three-piece Gibson assembly reaction that contained the following PCR products and transforming into ER475. (1) a 4.7 kb fragment containing the sequence upstream of *amyE*, the erythromycin resistance cassette, and the Phyperspank promoter amplified from ER300 genomic DNA using oZA005 and oZA021 (2) a 248 bp fragment ordered from IDT, oZA025 (3) a 1.487 kb fragment containing ubiquitin and the sequence downstream of *amyE* amplified from ER565 genomic DNA using oZA002 and oZA023.

**ER570** harboring amyE::Phyperspank-ubiquitin-ubiquitin-ubiquitin-TC::erm was constructed by performing a two-piece Gibson assembly reaction that contained the following PCR products and transforming into ER475. (1) a 5.02 kb fragment containing the sequence upstream of *amyE*, the erythromycin resistance cassette, the Phyperspank promoter, ubiquitin, and half of the sequence for the second ubiquitin were amplified from ER568 genomic DNA using oZA005 and oZA027 (2) a 1.657 kb fragment containing the second half of the sequence for the first ubiquitin, the second ubiquitin (including the FlAsH tag), and the sequence downstream of *amyE* were amplified from ER568 genomic DNA using oZA002 and oZA026.

**ER584** harboring amyE::Phyperspank-ubiquitin-TC-K::erm was constructed by performing a three-piece Gibson assembly reaction that contained the following PCR products and transforming into ER475. (1) a 2.888 kb fragment containing the sequence upstream of *amyE* and an erythromycin resistance cassette were amplified from ER565 genomic DNA using oZA066 and oZA069 (2) a 2.147 kb fragment containing the Phyperspank promoter and FlAsH-tagged ubiquitin with a terminal lysine residue were amplified from ER565 genomic DNA using oZA068 and oZA071 (3) a 1.222 kb fragment containing with FlAsH sequence with a terminal lysine residue and the sequence downstream of *amyE* were amplified from ER565 genomic DNA using oZA067 and oZA070.

**ER607** and **ER608**, harboring amyE::Phyperspank-ubiquitin-TC::erm and amyE::Phyperspank-mNeonGreen::erm, respectively, in a Δ*ponA*::kan background were generated by transforming ER478 with genomic DNA from ER565 and ER300 and selecting on LB + erm + kan.

**ER610** harboring amyE::Phyperspank-ubiquitin-TC-D::erm was constructed by performing a three-piece Gibson assembly reaction that contained the following PCR products and transforming into ER475. (1) a 2.888 kb fragment containing the sequence upstream of *amyE* and an erythromycin resistance cassette were amplified from ER565 genomic DNA using oZA066 and oZA069 (2) a 2.147 kb fragment containing the Phyperspank promoter and FlAsH-tagged ubiquitin with a terminal aspartic acid residue were amplified from ER565 genomic DNA using oZA068 and oZA073 (3) a 1.222 kb fragment containing with FlAsH sequence with a terminal aspartic acid residue and the sequence downstream of *amyE* were amplified from ER565 genomic DNA using oZA067 and oZA072.

**ER670** harboring amyE::Phyperspank-mNeonGreen::erm in a ΔlytCΔlytD ΔslrR::kan background was constructed by performing a three-piece Gibson assembly reaction that contained the following PCR products and transforming into ER669. (1) a 2.897 kb fragment containing the sequence upstream of *amyE* and an erythromycin resistance cassette were amplified from ER300 genomic DNA using oZA005 and oZA019 (2) a 1.899 kb fragment containing the Phyperspank promoter was amplified from ER300 genomic DNA using oZA003 and oZA020 (3) a 2.027 kb fragment containing mNeonGreen and the sequence downstream of *amyE* were amplified from ER300 genomic DNA using oZA051 and oZA054.

**ER672** harboring amyE::Phyperspank-mNeonGreen::erm in a ΔslrR::kan background was constructed by performing a three-piece Gibson assembly reaction that contained the following PCR products and transforming into ER671. (1) a 2.897 kb fragment containing the sequence upstream of *amyE* and an erythromycin resistance cassette were amplified from ER300 genomic DNA using oZA005 and oZA019 (2) a 1.899 kb fragment containing the Phyperspank promoter was amplified from ER300 genomic DNA using oZA003 and oZA020 (3) a 2.027 kb fragment containing mNeonGreen and the sequence downstream of *amyE* were amplified from ER300 genomic DNA using oZA051 and oZA054.

**ER728** harboring amyE::Phyperspank-ubiquitin-TC::erm in a ΔslrR::kan background was constructed by performing a three-piece Gibson assembly reaction that contained the following PCR products and transforming into ER671. (1) a 2.897 kb fragment containing the sequence upstream of *amyE* and an erythromycin resistance cassette were amplified from ER300 genomic DNA using oZA005 and oZA019 (2) a 1.899 kb fragment containing the Phyperspank promoter was amplified from ER300 genomic DNA using oZA003 and oZA020 (3) a 1.115 kb fragment containing the FlAsH-tagged ubiquitin and the sequence downstream of *amyE* were amplified from ER565 genomic DNA using oAE4 and oAE5.

**ER729** harboring amyE::Phyperspank-ubiquitin-TC::erm in a ΔsigD::kan background was constructed by performing a three-piece Gibson assembly reaction that contained the following PCR products and transforming into ER578. (1) a 2.897 kb fragment containing the sequence upstream of *amyE* and an erythromycin resistance cassette were amplified from ER300 genomic DNA using oZA005 and oZA019 (2) a 1.899 kb fragment containing the Phyperspank promoter was amplified from ER300 genomic DNA using oZA003 and oZA020 (3) a 1.115 kb fragment containing the FlAsH-tagged ubiquitin and the sequence downstream of *amyE* were amplified from ER565 genomic DNA using oAE4 and oAE5.

### Single-cell microfluidics permeability assays

Overnight cultures were diluted 100-fold into LB + 2 mM IPTG and incubated at 37 C for 2.5 hours. For the tetracysteine-tagged probes, 1 μM FlAsH-EDT_2_ was added to the back dilution one hour prior to imaging. Cultures were diluted 40-fold into a pre-warmed medium and loaded into a CellASIC B04A microfluidic flow cell (Millipore Sigma).

To ensure that the cells were growing at steady state, before the cells were imaged the cells were incubated for an additional 30 minutes in the microfluidics device within the microscope environmental chamber, which was pre-heated to 37 C. Before loading the cells into the imaging chamber of the flow cell, the chamber was primed with growth medium using the ONIX microfluidic perfusion platform. While imaging, fresh medium was perfused through the flow cell. The cell-trapping mechanism used by the microfluidic chips had no effect on the growth or morphology of cells, as compared with cells growing on agarose pads or liquid culture (43).

During phosphate buffer saline (PBS) incubation, the growth medium in the flow cell was exchanged with PBS (no divalent cation supplements) + 2 mM IPTG using the ONIX system. To perform membrane lysis, the growth medium in the flow cell was exchanged with LB + 5% N-lauroylsarcosine (Sigma Aldrich) + Alexa Fluor 647 Succinimidyl Ester (Invitrogen). The Alexa Fluor 647 dye allowed us to verify the switch to detergent. Two minutes after membrane lysis, the detergent in the flow cell was exchanged for LB, unless otherwise stated.

Images were acquired using a Nikon Ti-2 Eclipse inverted microscope and a Prime BSI sCMOS camera (Teledyne Photometrics). The intensity of cytoplasmic mNeonGreen and FlAsH-EDT_2_ were imaged using 480 nm excitation from an LED light source (Lumencor Aura Light Engine). mNeonGreen was imaged at 100% intensity, 50 ms exposure and FlAsH-EDT_2_ was imaged at 10% intensity, 50 ms exposure. Both were imaged at a one-minute frame rate for the first ten minutes and then a five-minute frame rate for the subsequent duration of the experiment. The Alexa Fluor 647 was imaged using 640 nm excitation at 10% intensity, 50 ms exposure.

### Agarose pad permeability assay

*B. subtilis* PY79 amyE::ermR-Phyperspank-mNeonGreen (ER300) was grown in LB in an shaking incubator at 30 C. Before microscopy, cultures were maintained in exponential phase for >10 population doubling times. Final OD was within 0.1-0.2 in all replicates. During microscopy, cells were sandwiched between a 1% agarose pad (made with LB or PBS) and glass coverslip. The agarose pad was 0.7 x 0.7 x 0.15 cm^3^. Before adding them to agarose pad (UltraPure Agarose, Invitrogen), bacteria were washed twice with pre-warmed LB or PBS. A 1 µL aliquot of bacterial culture with an OD600 of approximately 0.1 was directly deposited onto a 35 mm glass coverslip bottom dish (IBIDI), pre-assembled on the microscope and situated within a temperature-controlled stage-top incubator (at 30 C). The droplet was promptly covered with the agarose pad. Imaging began within 1.5-2.5 minutes after extracting cells from the shaking incubator.

Concentrated N-lauroylsarcosine solution in LB or PBS that had been pre-warmed to 30 C was added to top of the pad and allowed to diffuse in (final concentration 1% NLS). Time-lapse experiments were conducted on agarose pads (made with LB or PBS), with frame rate ranging from 60-180 s.

Imaging was performed using a Nikon Ti-E inverted microscope featuring both phase-contrast and epi-fluorescence capabilities, enhanced with a custom-designed module for SLIM (Spatial Light Interference Microscopy). The microscope setup comprises a temperature chamber (Stage Top Incubator, Okolab), a Nikon Plan Apo 100× NA 1.45 Ph3 objective lens, a solid-state light source (Spectra X, Lumencor Inc., Beaverton, OR), a multiband dichroic mirror (69002bs, Chroma Technology Corp., Bellows Falls, VT), and excitation (485/25) and emission (535/50) filters tailored for GFP imaging. We acquired epifluorescence images using a sCMOS camera (Orca Flash 4.0, Hamamatsu). Phase-contrast and quantitative phase images were captured on a separate CMOS camera (DCC3260M, Thorlabs). The methodology for microscopy and subsequent image analysis has been detailed in prior work (30).

In brief, quantitative phase images from four distinct phase-contrast images were generated, each incorporating an additional π/2 phase shift of the non-scattered light, facilitated by a spatial light modulator. MATLAB-based Morphometrics package was employed to extract cell dimensions from phase-contrast images. Surface area and volume were derived from corrected cell contours with an assumption of cylindrical symmetry around the cell axis. Cell mass was calculated by integrating the phase shift of individual cells.

### Image analysis

Image analysis was performed using custom MATLAB (Mathworks, Natick, MA, USA) scripts. Cell boundaries were tracked using phase contrast images. Phase contrast images were aligned to fluorescence images. The cellular fluorescence trace was obtained by measuring the averaged fluorescence intensity of each tracked cell.

The background fluorescence was subtracted from each cellular fluorescence trace. The fluorescence from the frames during lysis were interpolated. The cellular trace was corrected for photobleaching and then normalized to the final pre-lysis frame.

To perform the photobleaching controls (**Fig. S9, S10**), cells were imaged at one frame per minute during the initial seven minutes of the experiment prior to cell lysis. After membrane lysis, the cells were imaged at frame rates that were much more rapid that protein efflux.

### Photobleaching correction

Population-averaged fluorescence traces were fit to a single exponential:

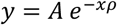

where y = normalized fluorescence, A is the initial post-lysis normalized fluorescence value, ρ = the photobleaching constant, and x is the frame number (Figure S9, S10).

For mNeonGreen, the mean background value is subtracted from the mean fluorescence of each cell and normalized by the fluorescence of the final pre-lysis frame.

For FlAsH, the mean background value and mean autofluorescence is subtracted from the mean fluorescence of each cell. The mean autofluorescence was obtained by measuring the fluorescence of uninduced cells incubated with FlAsH-EDT_2_.

To correct for photobleaching, the expected loss of fluorescence due to photobleaching was calculated,

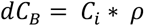

where C_B_ = bleached fluorophore, C_i_ = measured fluorescence in the *i*th frame, and π = photobleaching constant (obtained from the photobleaching control). The total loss of fluorescence (dC_T_) in each frame is adjusted to account for the expected loss of fluorescence due to photobleaching,

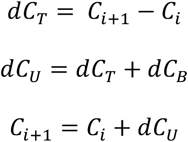

where dC_U_ is the change in unbleached fluorophore.

### Lag time and efflux calculation

Normalized fluorescence traces were interpolated and smoothed, and then the first derivative was calculated by finding the slope of each trace along a moving window. The efflux rate was defined as the minimum of the first derivative of the normalized fluorescence. To calculate lag time, we calculated the time at which efflux was maximum, extrapolated a line with the same slope as the efflux rate, and found the time at which this line intercepted 1, which we defined as the lag time lag time.

### Phospholipase assay

Overnight cultures were diluted 100-fold into LB and incubated at 37 C for 2.5 hours. The back dilutions were further diluted 100-fold into media in a 5 mL sample tube. 2 μg/mL propidium iodide was added to the media. At 45 seconds after beginning a cytometry run, 2 μg/mL phospholipase A2 was added through a port in the sample tube. To permeabilize the cells, cells were washed twice with PBS and then added to the sample tube. The experiment was performed using a Cytek Aurora flow cytometer on the low flow rate setting. The flow cell was cleaned between each experimental run. Propidium iodide positive and negative gates were manually drawn using SpectroFlo software, exported to .fcs files, and analyzed using a custom python script. The python script utilizes the FlowKit package (48).

### FDAA release assay

Overnight cultures were diluted 100-fold into LB and incubated at 37 C for 2.5 hours. One hour prior to the end of the back dilution, 100 μM of NADA-green (Tocris), a fluorescent D-amino acid (FDAA), was added to the culture. For the dead stain, 2 μg/mL propidium iodide was added to the media. Only propidium-iodide negative cells were used in the analysis. At the end of the back dilution, the culture was washed twice with LB. The back dilutions were further diluted 100-fold into media in a 5 mL sample tube. One-minute runs were performed over a series of time points (t = 0, 1, 2, 2.5, and 4.5 hours). The NADA fluorescence (B2 channel) was normalized to t = 0, interpolated, and fit to a single exponential to calculate the time scale of hydrolysis.

### Charged surface representation

The PDB structures for phospholipase (1POC) and mNeonGreen (5LTR) were downloaded from the Protein Data Bank (49, 50). The PDB structure for mUbq was generating using ColabFold and the AlphaFold2-mmseqs model. Based on these PDB structures, the charged surface representation were generated using the APBS Electrostatics plugin in Pymol (51).

### Molecular properties

The isoelectric point and molecular weights for each protein were calculated by ExPasy using the protein sequence (52). The charge was calculated by Prot Pi using the protein sequence. The radius of translation was calculated by HullRad using the PDB structures (53).

### Optical density assay

Overnight cultures were diluted 100-fold in LB and incubated at 37 C for 2 (*E. coli*, ER005) and 2.5 hours (*B. subtilis*, ER475). The OD600 was measured using a GENESYS 140 Vis Spectrophotometer (Thermo Scientific). Optical density was measured again after the cells were resuspend in LB + 5% NLS. The fractional change was calculated by the following:

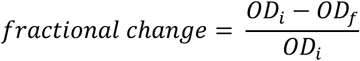

Where OD_i_ is the pre-lysis optical density and OD_f_ is the post-lysis optical density.

### Agarose pad imaging of FlAsH-tagged ubiqiutin strains

Overnight cultures were diluted 100-fold in LB (uninduced) or LB + 2 mM IPTG (induced) and incubated at 37 C for 2.5 hours. One hour prior to imaging, 100 mM TAMRA-based fluorescent D-amino acid was added to the induced cultures and 1 μM FlAsH-EDT_2_ was added to all the cultures. The cultures were washed with LB, and then 200 μL of the induced and uninduced cultures were mixed together in a new tube. The mixed culture was added to a 1.5% (low melting) agar pad and imaged at 100X magnification in phase, GFP (480 nm excitation, 10% intensity, 50 ms exposure), and RFP (560 excitation, 10% intensity, 100 ms exposure).

### PAGE electrophoresis of ubiquitin probes

Overnight cultures were diluted 100-fold into LB (+ 2 mM IPTG if induced) and incubated at 37 C for 2.5 hours in a final volume of 22 mL. FlAsH-EDT_2_ was added one hour prior to the end of the back dilution for a final concentration of 1 μM. Cultures were kept away from light to prevent photobleaching. Cells were pelleted and incubated with lysis buffer at 37 C for 10 minutes. After adding Lammeli buffer, samples were boiled at 80 C for 5 minutes. The samples and ladder (BioRad Precision Plus All Blue Prestained Protein Standards) were loaded onto a 12.5% SDS-PAGE gel and run at 150 V. The gel was imaged on an Amersham Typhoon imager (the FlAsH-EDT_2_ in the Cy3 channel, the ladder in the Cy5 channel). Images were merged and false colored using Fiji (54).

### Combinatorial staining of strains expressing charge-variants of mono-ubiquitins

Overnight cultures of amyE::Phyperspank-ubiquitin-TC::erm (ER565, ER584, ER610), were diluted 100-fold into LB + 2 mM IPTG (+ 1 μM FlAsH-EDT2 and 100 μM of TADA or HADA fluorescent D-amino acids, one hour prior to imaging) and incubated at 37 C until the cells were in mid-exponential phase. Equal volumes of back diluted culture were aliquoted together. The combined sample was diluted 40-fold in LB and 200 μL of this dilution was added to the microfluidics device and cells were loaded. Before adding the cells to the device, the chamber was primed with growth medium using the ONIX microfluidic perfusion platform (CellASIC).

### Protoplast assay

Overnight cultures of ER300 (amyE::ermR-Phyperspank-mNeonGreen) were diluted 1:100 fold in LB + 2 mM IPTG. At the end of the back dilution, washed twice with 1xSMM media (500 mM sucrose, 20 mM maleic acid, 20 mM magnesium chloride hexahydrate, pH 6.5) and incubated for 1 hour with 0.5 mg/mL lysozyme with gentle rocking.

### Analysis of *ΔsigD* fluorescence

The permeability assay was set up as previously described, but due to chaining the cells could not load properly into the microfluidics device. To measure cellular fluorescence, in Fiji intensity was measured from points marked along the cell chains in each frame. To measure background fluorescence, intensity was measured from points marked away from cell chains. To calculate the normalized uncorrected fluorescence, the background and uninduced autofluorescence (based on WT measurements) were subtracted from the average intensity value for all the points in each frame. This value was normalized to the pre-lysis frame. For WT, a similar calculation was performed except with intensities from cells tracked via a custom MATLAB program.

### Derivation of the scaling rule given by Eq. 1

The diffusion constant, *D*, defines the flux of particles, *J,* diffusing through a surface given a spatial gradient ∂C/∂x in the direction normal the surface:

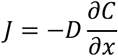

The rate at which concentration in the sacculus changes is then

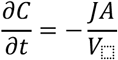

where *A* is the surface area of the cell covered by pores and *V*_⬚_is the volume of the cell. Combining these equations, we obtain

The time scale at which particles diffuse at the length scales in question is much faster than the time scale at which concentration decreases (minutes). In other words, once particles leave the sacculus they will rapidly diffuse away, and concentration rapidly equilibrates within the sacculus. Therefore, we may assume that the concentration gradient across the thickness of the cell wall is equal to the concentration of particles in the sacculus, *C*_in_, divided by cell wall thickness, δ,

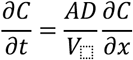

Therefore, the time scale that the concentration inside the sacculus, initially at *C*_in_, decreases by a factor of two is given by Eq. 1

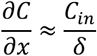

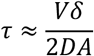

Using *D* = 1μm^2^/s (55), δ = 0.032 μm (19) *V* = πR^2^l, where R = 0.4 μm and l = 4 μm, we find that τ ≈3 ms if the entire cell surface were freely permeable to the particles. On the other hand, given that the experimental time scale for protein efflux is τ ≈ 2 *min*, the area of the cell surface that is expected to be covered with pores is *A* ≈ 8.4 × 10^-5^ μm^2^. If the cross-sectional area of a single mono-ubiquitin protein is equal to *A*_)45_ ≈ πR^2^ = 1.0 × 10^-5^ μm^2^ where R is the radius of translation (Table 1). According to this dimensional order-of-magnitude calculation, there only needs to be 26 mono-ubiquitin-sized pores for efflux to occur at experimental time scales.

An alternative to the discrete-pore model for the relatively slow efflux of mono-ubiquitin is that the probe diffuses through the cell wall at all points, but at a slower rate than if were diffusing freely. However, there are two related arguments against this explanation. First, slow diffusion of the probe independent of discrete pores would be analogous to the mechanism by which polyacrylamide slows proteins migration during gel electrophoresis: in this case, slower diffusion of larger proteins occurs because there are many pores with a continuous distribution of sizes. If this were the case for the cell wall, we would expect bi-ubiquitin and mNeonGreen efflux to occur more slowly, but not be completely blocked by the wall. Second, if the cell wall meaningfully slowed diffusion, this would cause a delay in the onset of efflux, followed by rapid disappearance of the probe (as during electrophoresis, it takes time for protein to migrate a certain distance, but it all arrives at the same time). In fact, we observe evidence of a slight delay in protein efflux on the order of seconds, which could be due to this effect, but it is negligible compared to the time scale of protein efflux, arguing for a efflux through discrete pores.

### Computational percolation model

Proteins (particles) inside the cell are represented as random walkers on a two-dimensional square lattice with lattice spacing (site size) *a*. The cytoplasm is treated as a region of free diffusion with no particle-particle interactions, while the cell wall is modeled as a boundary in which a fraction, *p*, of sites are occupied by obstacles that block movement (reflecting boundary conditions). The survival (or retention) probability (plotted in Fig. 3C) is defined as the fraction of particles that remain inside the cell up to time *t*. A particle is considered to have escaped once it crosses the finite cell wall of width δ, at which point the exit time is recorded for computation of the survival probability.

The simulated geometry is illustrated in **Fig. 3A**. The bacterium is represented as a two-dimensional cross-section taken perpendicular to the long axis of the rod-shaped cell. Along the *x*-axis, the cell wall defines the left and right boundaries of the domain, while in the *y*-direction the cell extends much farther than its width. To reflect this, periodic boundary conditions were imposed in the *y* direction. The particles were initialized anywhere within the interior with uniform probability. The choice of cell height (table S2) ensured that within the simulated time window the particle was unlikely to traverse the full height, thereby avoiding repeated sampling of the same regions and providing a closer approximation to the physical system. For additional simplification, a reflecting boundary was placed along the central axis of the cell, rendering the system symmetric about the vertical axis. For various possible outcomes of simulated trajectories, see Fig. S6 which also showcases the dependence on wall density p.

To allow finer resolution of particle sizes, we refined the lattice while scaling particle dimensions consistently, ensuring that the effective geometry of the system remained unchanged. In the basal model, the lattice constant *a* (site size) corresponds to the characteristic pore diameter of a randomly cross-linked peptidoglycan. In this case, each site was subdivided into a 4 × 4 block of smaller sites (*a*^7^ = *a*/4), allowing the simulation of particles of size 3.6 and 4.5 nm with a discrete representation, *a*^’^ = 0.9*nm*. Here, we assumed that the characteristic pore size is the same as the smaller particle (3.6nm). To prepare different random realizations of the wall before simulation, each block of size 3.6 x 3.6 nm within the wall was assigned a state of occupied (value 1) or unoccupied (value 0) by choosing a uniform random number and checking whether it is less than *p*.

Simulations were performed using a random walk approach by randomly selecting (with uniform probability) a direction to move a protein by a constant step size Δl = 1 *nm*. The move was accepted if none of the squares which would be covered in the new position were occupied, and otherwise the move was rejected, with the time incremented by 1 time step Δ*t* in either case. Simulations were performed with experimentally derived parameters as indicated in **Table S2**.

The critical threshold *p*\* for a network of finite thickness δ was defined as the *p* at which 95% percent of particles, on average, are still trapped inside the cell after 1 minute of starting their motion (Fig.3B). The values are *p*\*=0.485 and *p*\*=0.75 for particles of size 3.6 nm and 4.5 nm, respectively.

### Strains

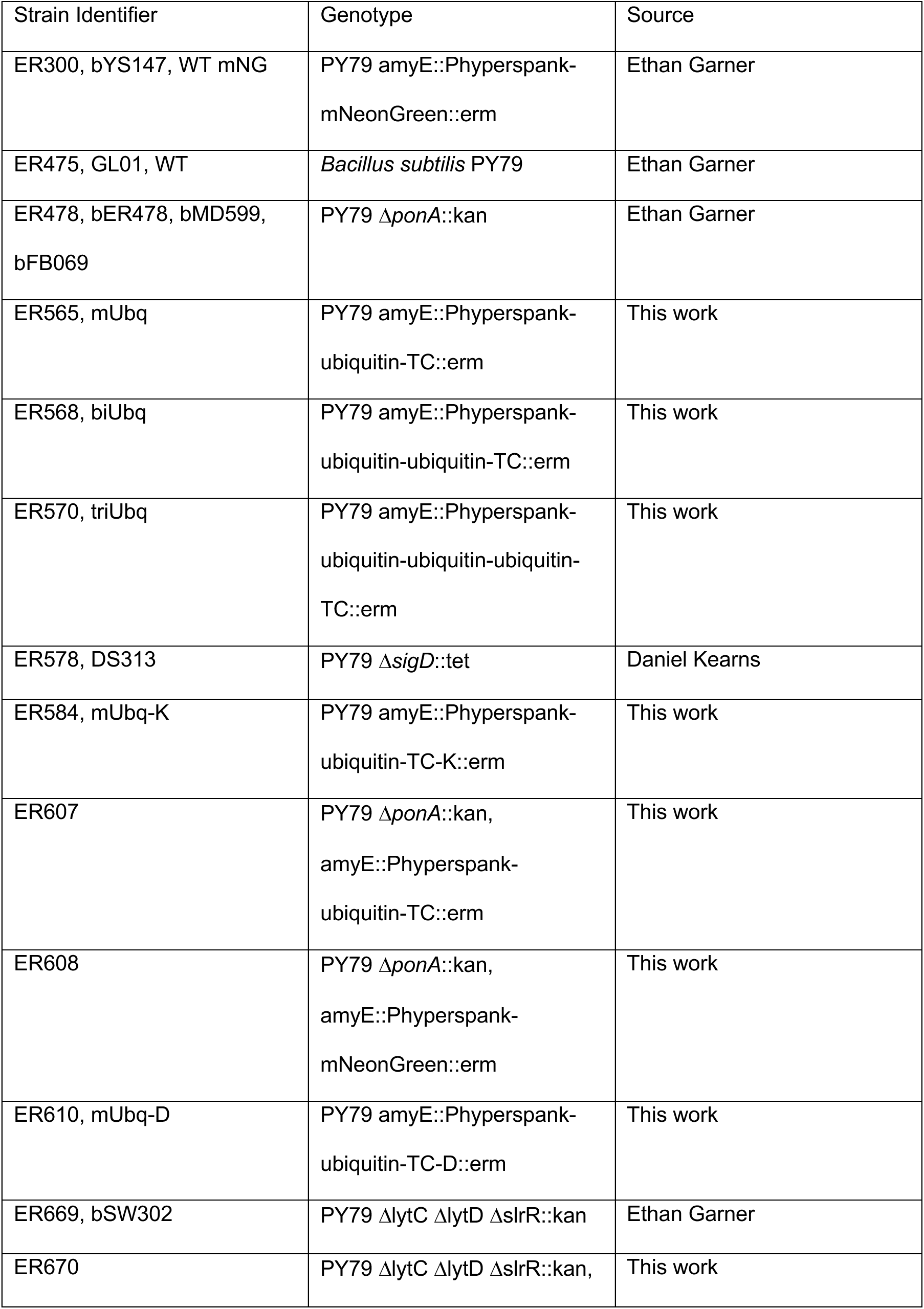
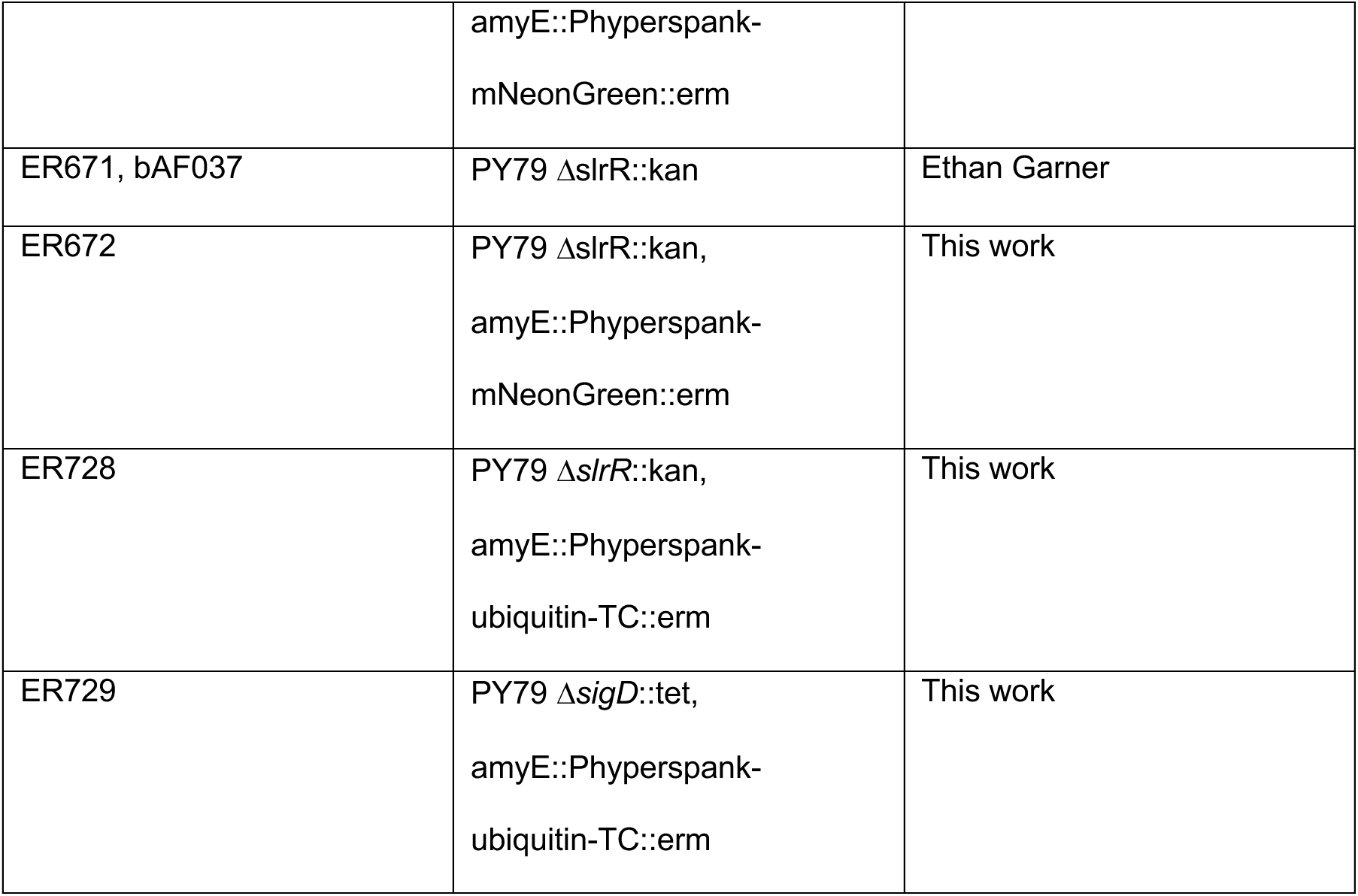

### Primers

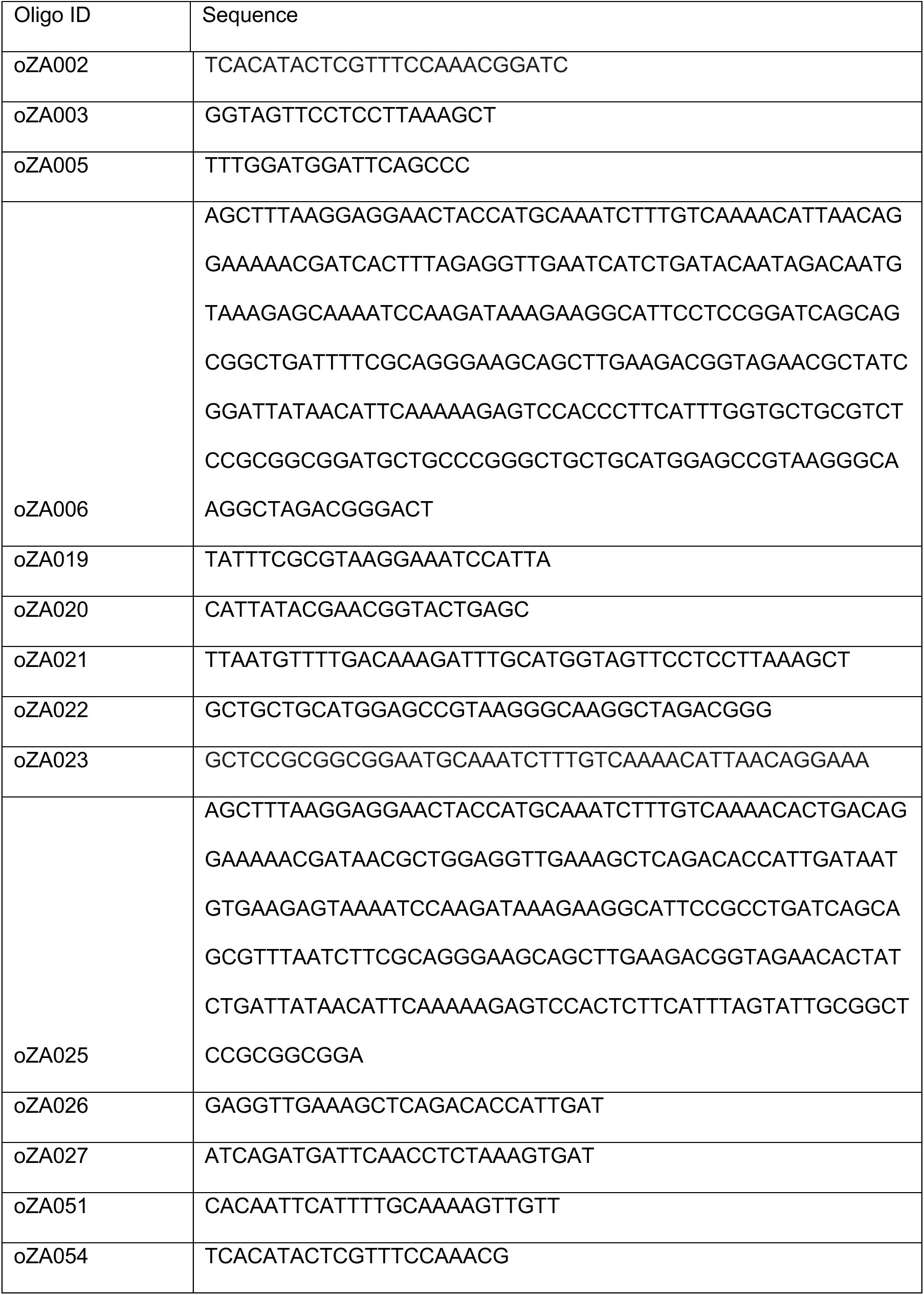
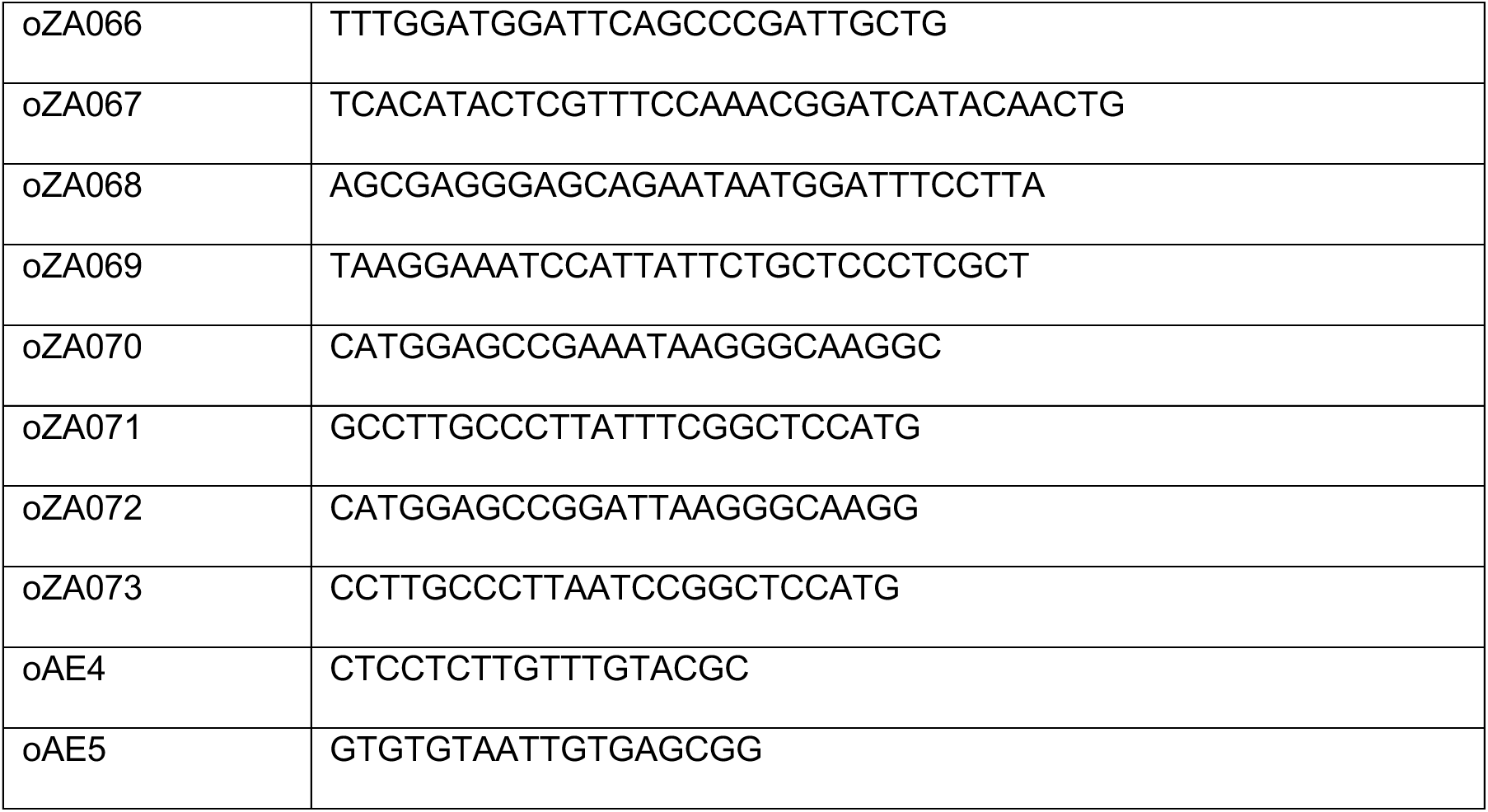

### Reagents

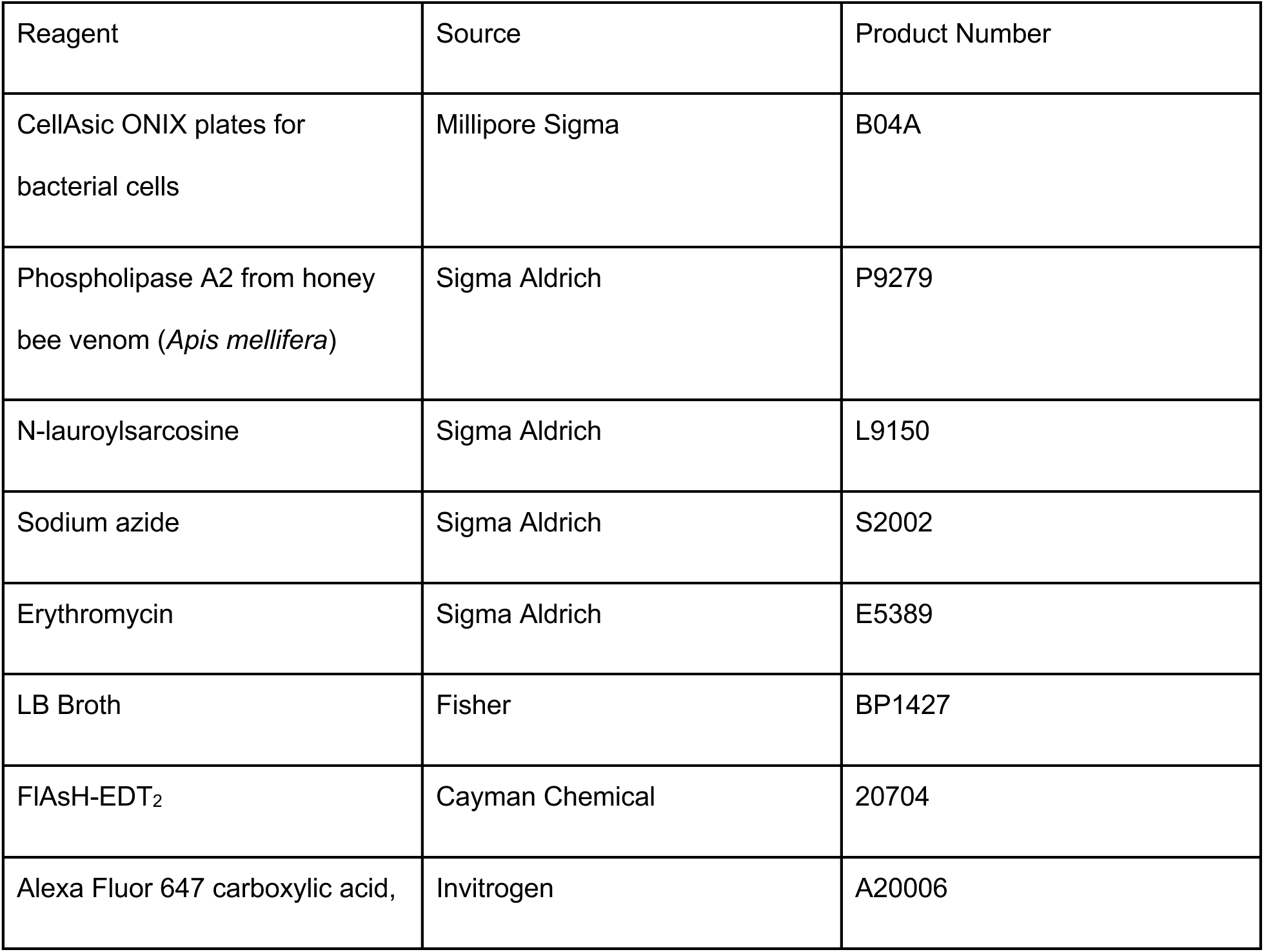
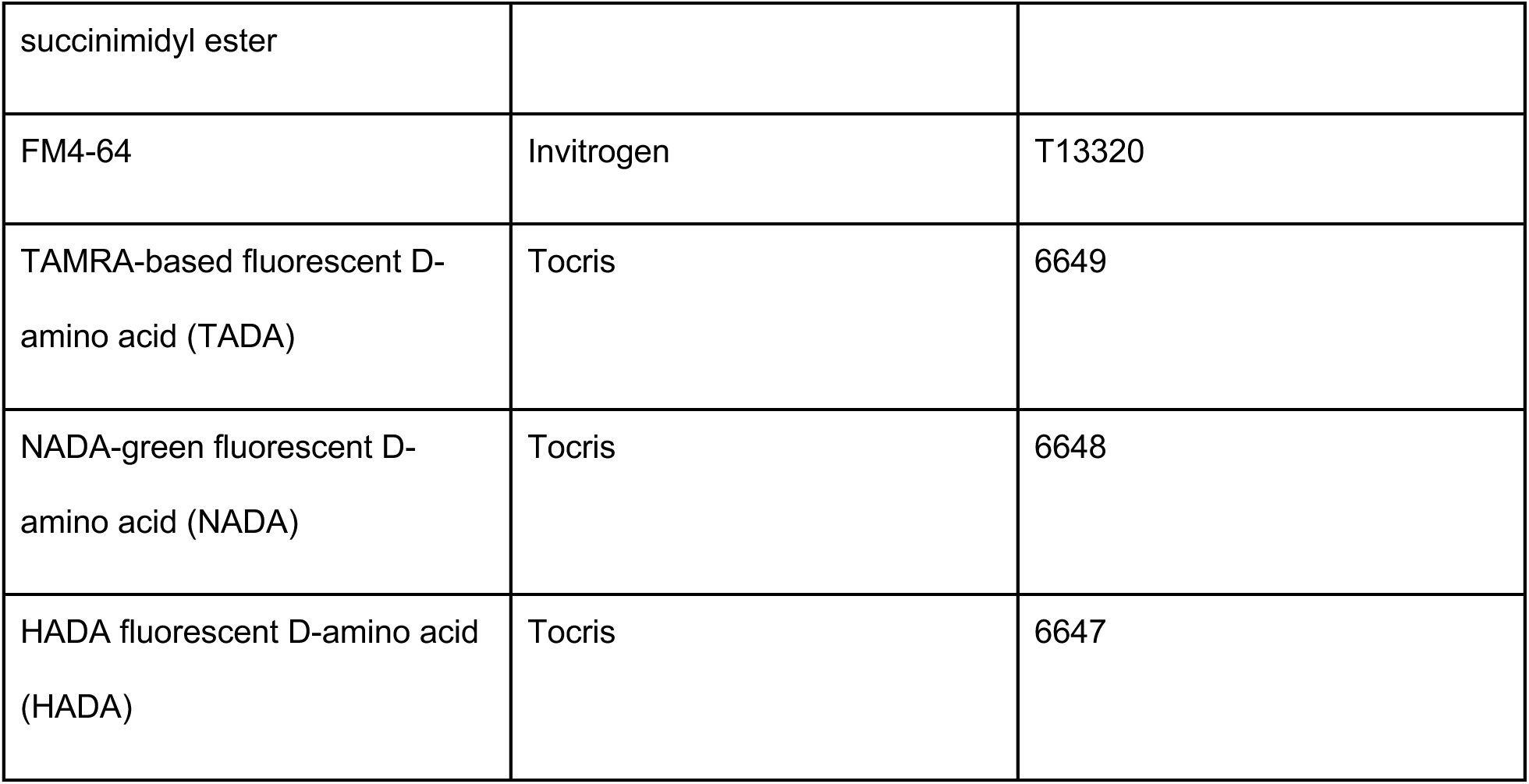

## Acknowledgments

The authors thank Ethan Garner, Sean Wilson, Dan Kearns, and Sadie Ruesewald for insightful discussions and bacterial strains, as well as Victor Leon and Harsh Srivastava for technical assistance. ER and ZA were supported by NIH Grant R35GM143057. ZA was additionally supported by NIH Training Grant T32GM132037. GMH and DS were supported by NIH Grant R35GM138312.

---

**Figure S1.**
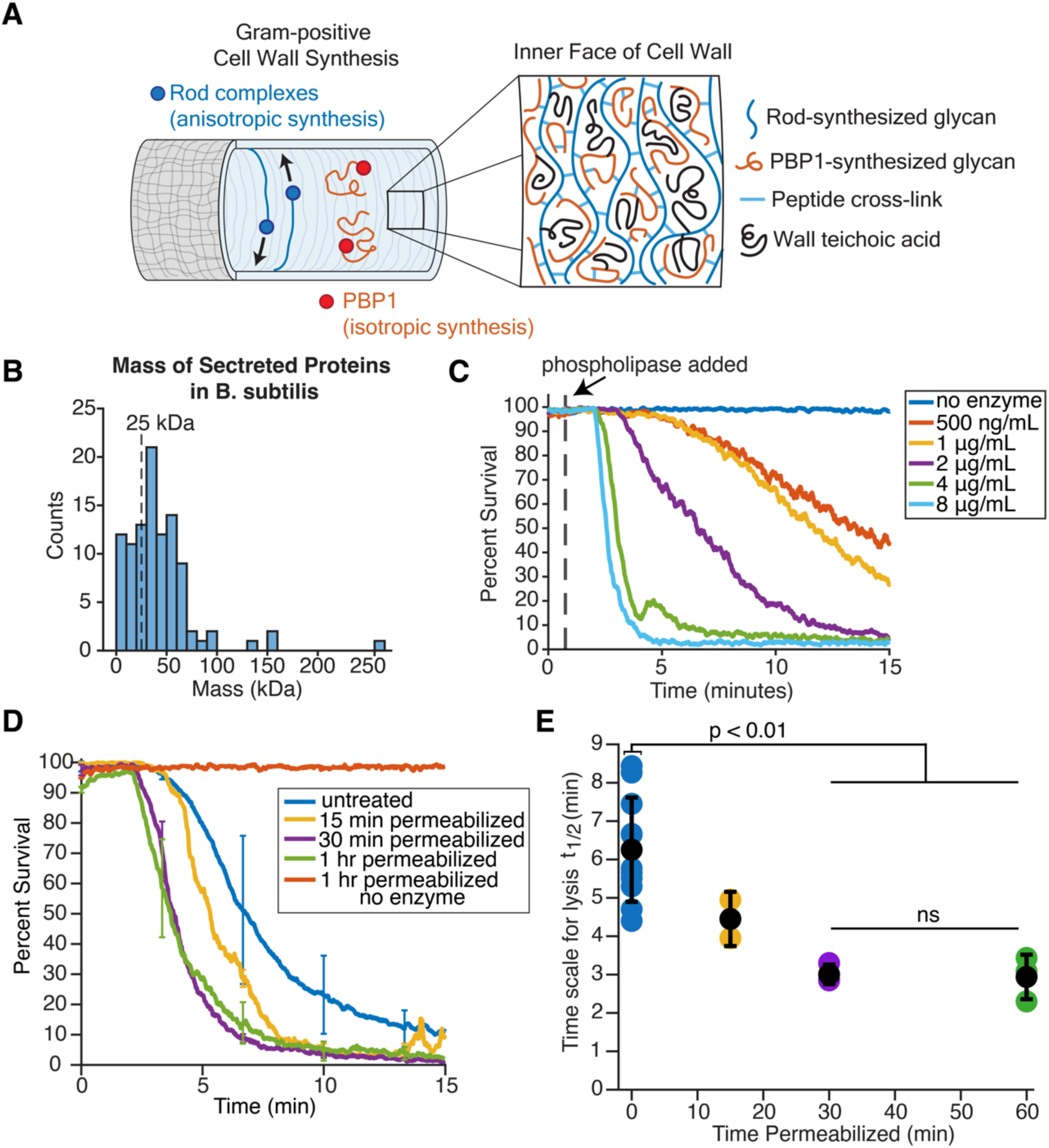
The cell wall acts as a barrier against phospholipase. (*A*) Diagram of the two modes of peptidoglycan synthesis. The Rod complex (blue) synthesizes peptidoglycan oriented along the circumference of the cell. PBP1 (red, encoded by *ponA*) synthesizes peptidoglycan isotropically. Inset: a diagram of the inner face of the cell wall with oriented Rod complex-synthesized strands (blue), non-oriented PBP1-synthesized strands (red), peptidoglycan cross-links (light blue) and wall teichoic acids (black). (*B*) Distribution of protein masses for secreted proteins in *B. subtilis* 168. Data obtained from the UniProt database after filtering for proteins with the ‘secreted’ label and without the ‘membrane’ label in the subcellular localization field (23). (*C*) Mean percent survival of cells treated with varying phospholipase concentrations over time. Phospholipase was added at t = 45 seconds, as indicated by the dashed line. Each experiment was performed once. *(D)* Mean percent survival of PBS permeabilized cells over time. Cells were incubated in PBS for 0, 15, 30, or 60 minutes prior to the experiment. Cells were treated with 2 μg/mL phospholipase at t = 45 seconds. Data are presented as mean ± standard deviation (SD). Shown here for comparison, the untreated and 30-minute permeabilized data from Fig. 1F*. N* = 11, 2, 3, 3, 1 replicates. *(E)* t_1/2_ (time it takes to reach 50% survival) versus time spent incubating in PBS. Colored dots represent the t_1/2_ values for individual experiments. The black dots indicate mean t_1/2_ values. The error bars are the standard deviation. Statistical significance calculated using Student’s *t*-test.

**Figure S2.**
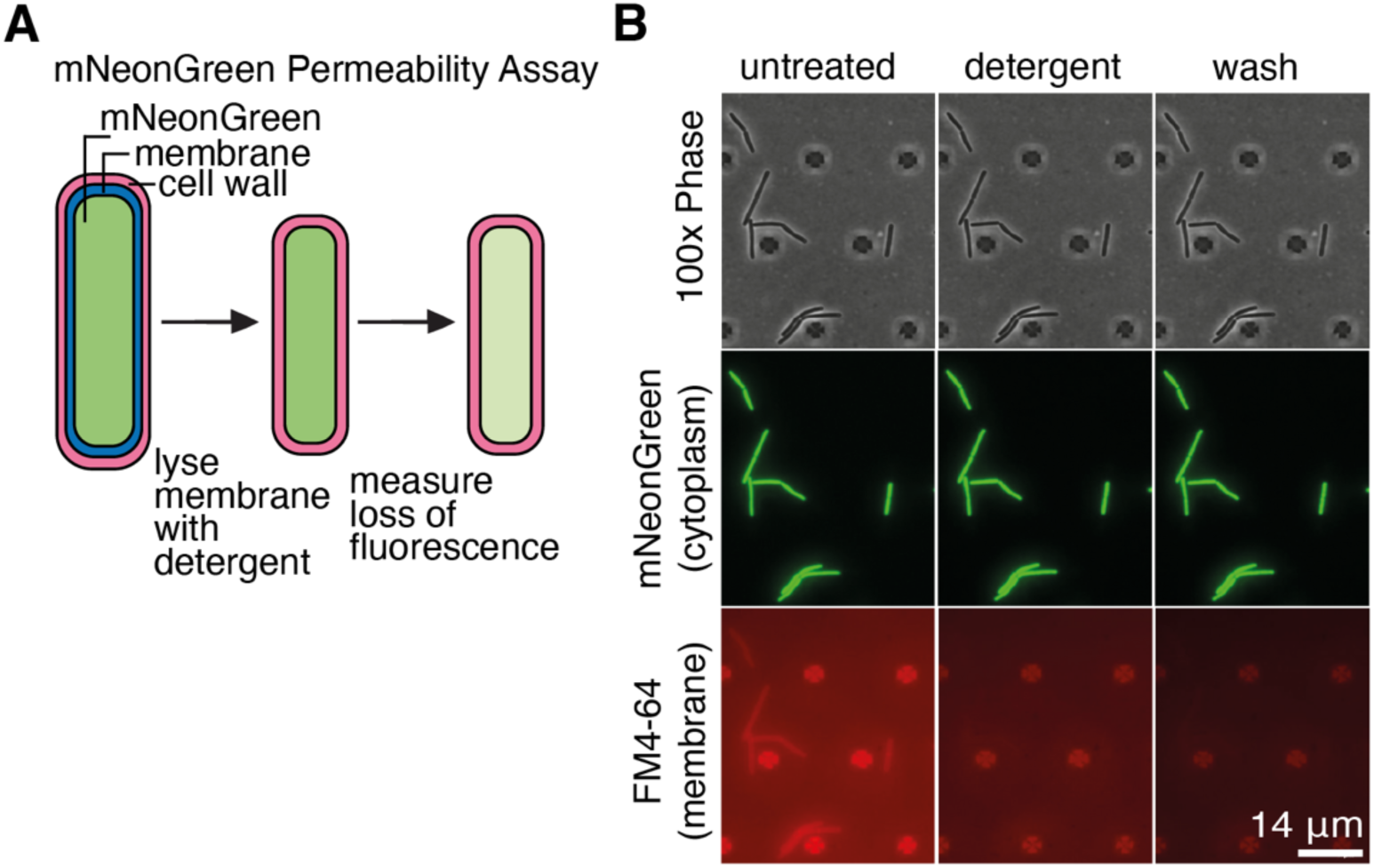
The cell wall sacculus can be isolated by treating cells with detergent. (*A*) Diagram illustrating the membrane lysis assay. Bacterial cells expressing cytoplasmic mNeonGreen (green) are subjected to detergent treatment to disrupt the membrane and kill the cell. Although the membrane undergoes lysis, the cell wall remains intact. After membrane lysis, loss of fluorescence is measured as an indicator of fluorescent protein diffusion through the cell wall. *(B)* Micrographs of *Bacillus subtilis* PY79 trapped in a microfluidics device. The top row shows the cells in 100X Phase, the second row shows cytoplasmic mNeonGreen, and the bottom row shows the plasma membrane stained with FM4-64. In the first column, cells are growing in rich media. In the second column, cells were treated with detergent (LB + 5% N-lauroylsarcosine). The plasma membrane ruptured but the cell shape (and ergo the cell wall) remained intact. In the third column, the cells were washed with LB to remove the detergent, but the plasma membrane does not re-assemble. Scale bar, 14 μm.

**Figure S3.**
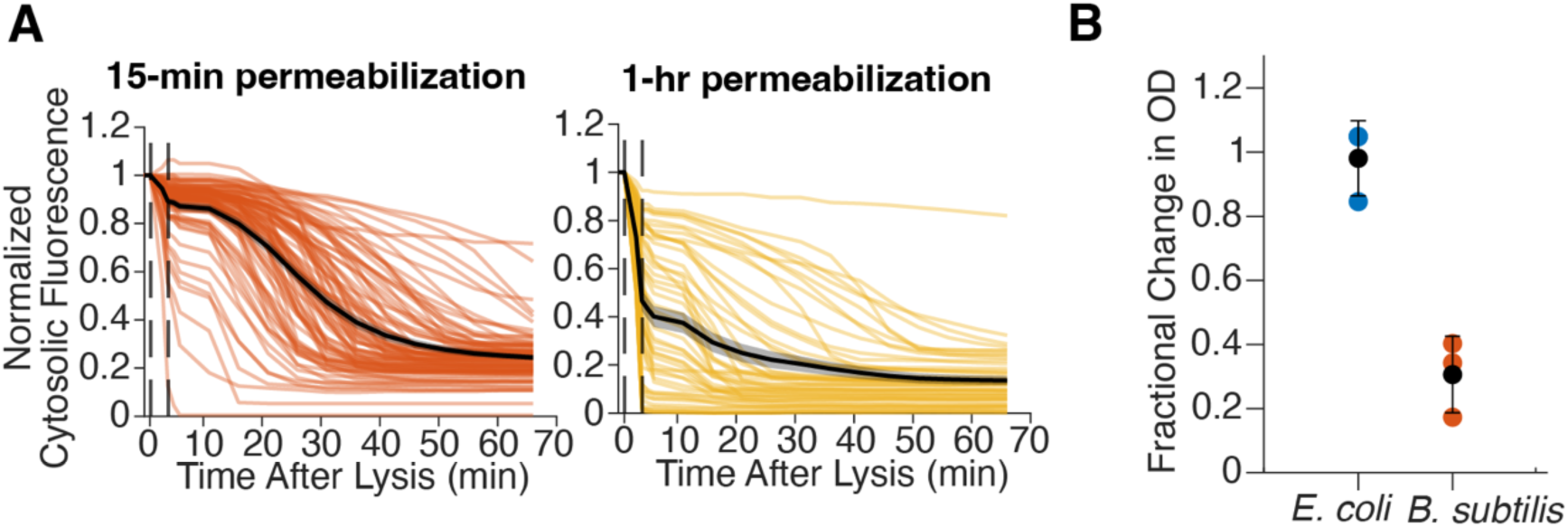
mNeonGreen efflux is positively correlated with the duration of autolysis. (*A*) Normalized fluorescence traces of the PBS-incubated cells shown in Fig. 2E. The black line indicates the mean normalized trace. The shaded region indicates the standard error of the mean. The first dashed black line indicates detergent perfusion. The second dashed black line indicates the wash. *(B)* Fold change in OD600 of *E.coli* and *B. subtilis* after resuspension in detergent (LB + 5% N-lauroylsarcosine). *N* = 3 replicates.

**Figure S4.**
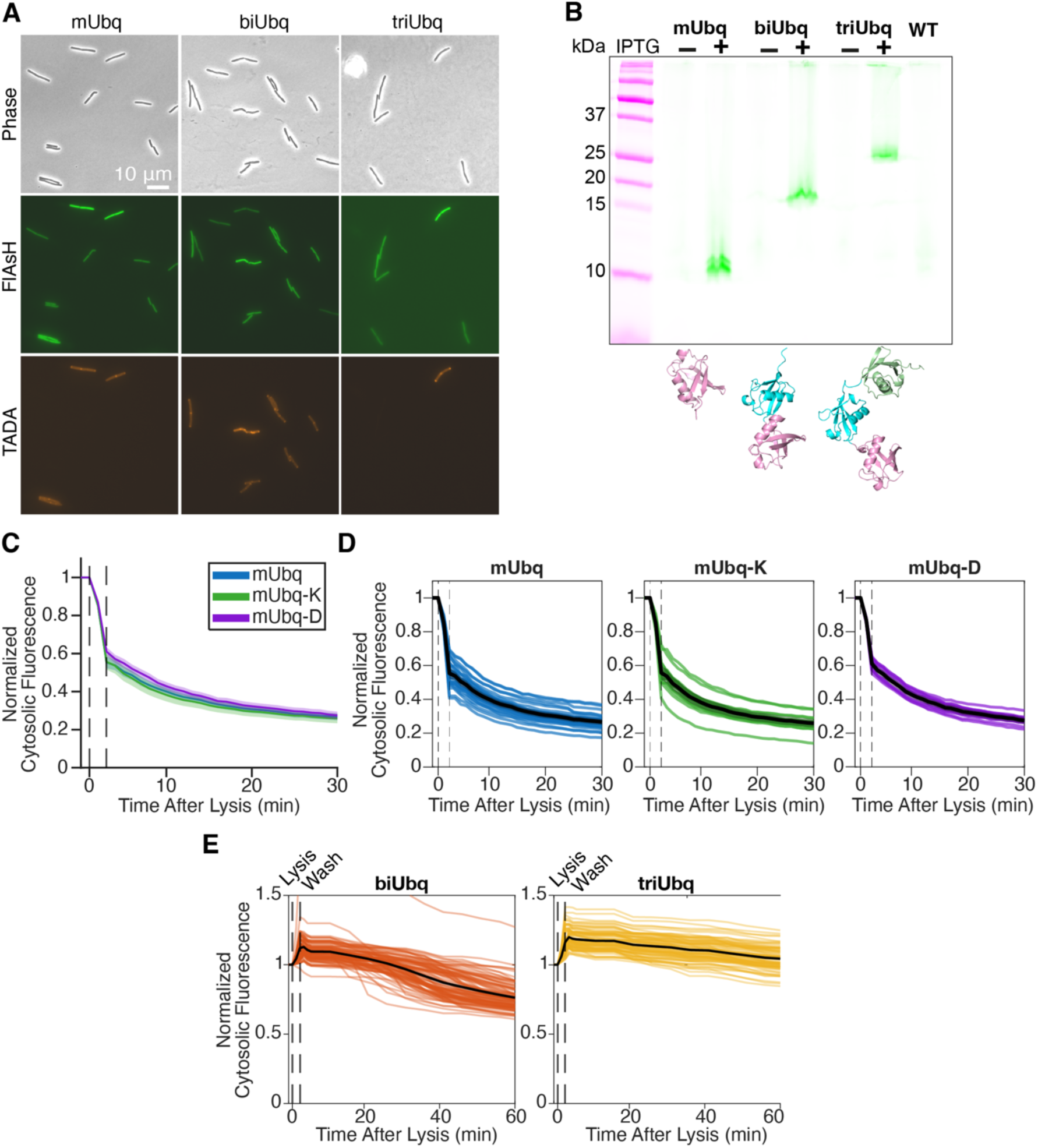
Chained ubiquitins do not diffuse through the cell wall and changing the isoelectric point of mono-ubiquitin (mUbq) does not alter diffusion. *(A)* Induced (TADA labeled) and uninduced (non-labeled) strains expressing mUbq (*left*), biUbq (*middle*), or triUbq (*right*) were incubated with 1 μM of FlAsH and added to an agarose pad. The top panel shows the cells in phase, the middle shows the fluorescence from the ligand, and the bottom panel shows the induced TADA-labeled cells. (*B*) Wild-type (WT) PY79 and mUbq, biUbq, and triUbq (uninduced and induced) were grown to exponential phase and incubated with 1 μM FlAsH. Cell lysates were run on a 12.5% SDS-PAGE gel. The fluorescent bands indicate protein bound to the FlAsH ligand, which are 10, 18, and 25 kDa for mUbq, biUbq, and triUbq, respectively. At the bottom of each lane for the ubiquitin probes are the corresponding AlphaFold structures. (*C*) The mean normalized fluorescence traces of mUbq (blue), mUbq-K (green), and mUbq-D (purple) expressing strains over time. The isoelectric point, as calculated by Expasy, are 5.74, 6.55, and 5.28, respectively (52). The first dashed line indicates the perfusion of detergent, the second dashed line indicates the wash. Cells were imaged at 10% intensity at a one-minute frame rate and corrected for photobleaching. The standard error of the mean is indicated by the shaded region. *n* = 26, 15, 10 cells across *N* = 3 technical replicates for mUbq, mUbq-K, and mUbq-D, respectively. (*D)* The individual traces for mUbq (blue), mUbq-K (green), and mUbq-D (purple) shown in *C*. In black, the mean of the cell traces. *(E)* Each individual trace represents the corrected, normalized fluorescence from a single cell expressing biUbq (orange, left) or triUbq (yellow, right). In black, the mean of the cell traces. *n* = 96, 79 cells across *N* = 2, 2 experiments for biUbq and triUbq, respectively.

**Figure S5.**
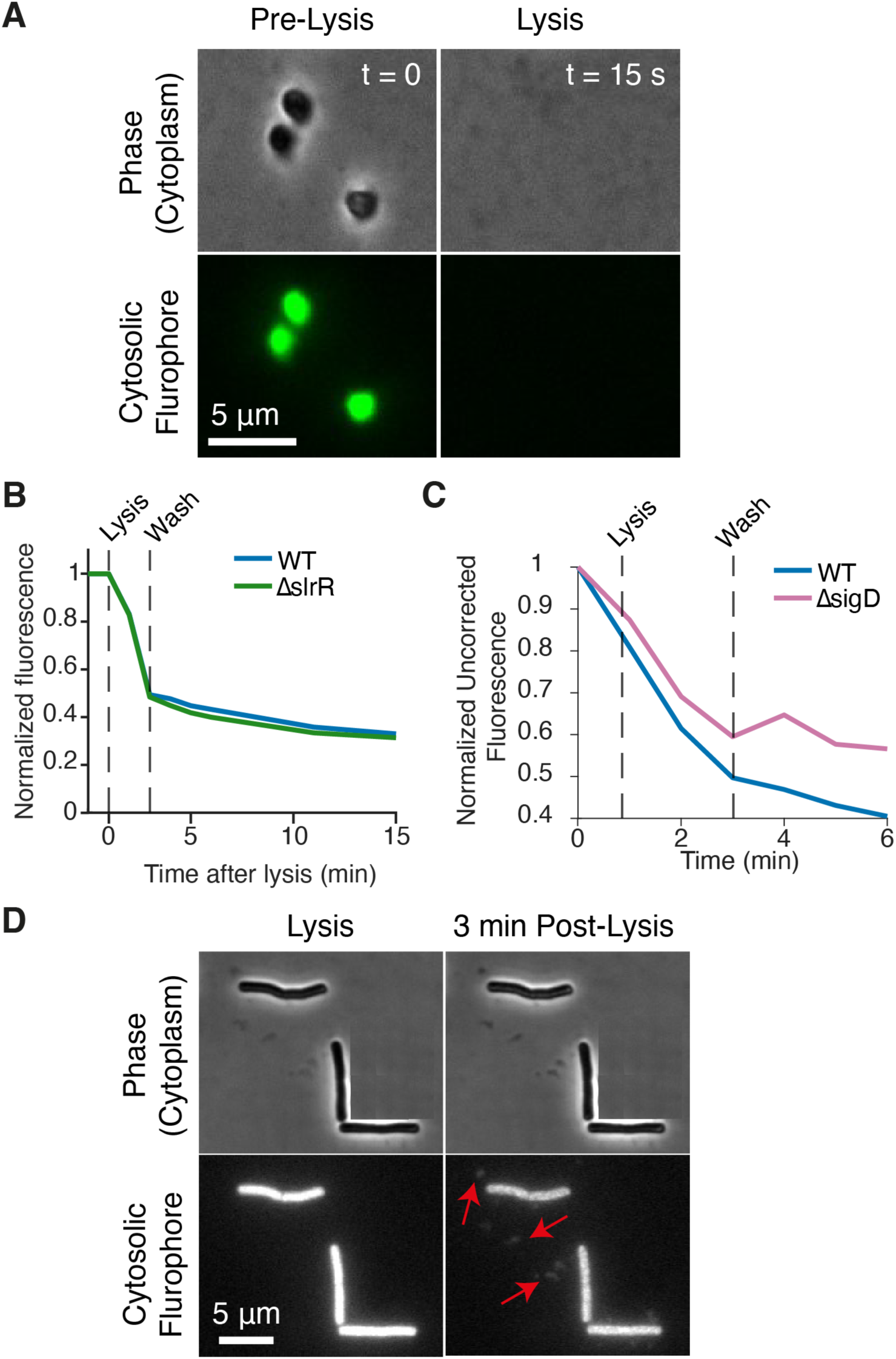
Motility mutants show no change in mono-ubiquitin (mUbq) efflux. (*A*) Micrographs of protoplasts expressing mNeonGreen before (pre-lysis) and 15 seconds after detergent perfusion. (*B*) Mean (left) and individual (right) normalized fluorescence traces of *ΔslrR* mUbq (green). Shown here for comparison, the wild-type (WT) mUbq data (blue) from Fig. 2G. Left, shaded region is the standard error of the mean. Right, the black trace is the mean normalized fluorescence. *n* = 176 cells across *N* = 3 technical replicates. (*C*) Mean normalized uncorrected fluorescence traces of *ΔsigD* mUbq (pink). Shown here for comparison, the mean normalized uncorrected fluorescence of WT mUbq data (blue) from Fig. 2G. *N* = 1 experiments, *n* = 10+ regions per frame. (*D*) Micrographs of protoplasts expressing mono-ubiquitin before (pre-lysis) and three minutes after detergent perfusion (3 min post-lysis). Red arrows indicate the fluorescent mono-ubiquitin aggregates outside of sacculi.

**Figure S6.**
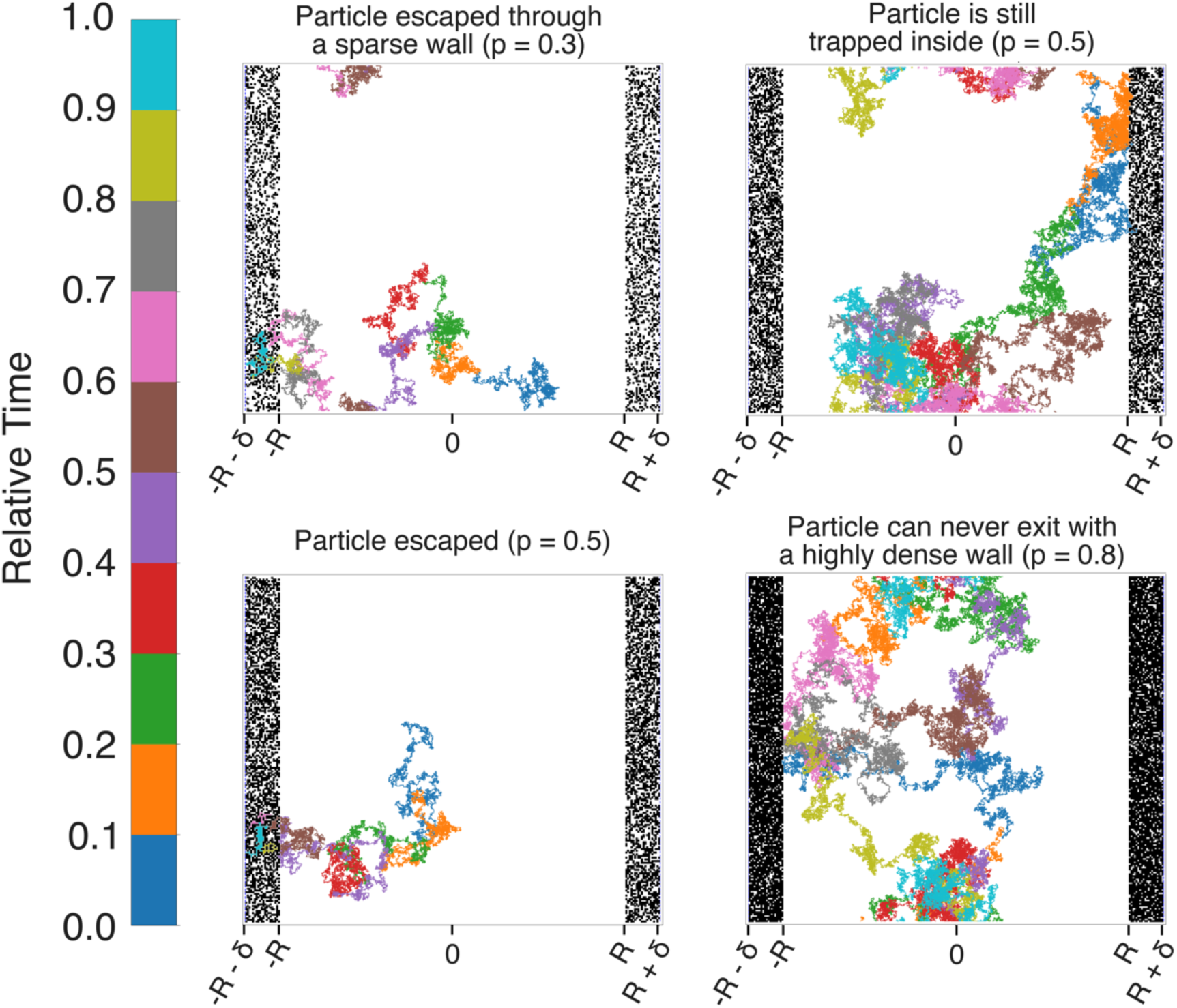
Simulated trajectory examples for different wall densities. Shown here are different possible outcomes with choice of parameters in Table S2. (A) a particle enclosed by a very sparse wall (p=0.3) will always be able to escape. (B) For more dense walls (p=0.5), a particle may still be trapped up to the maximum simulated time. (C) Particle escaped very quickly in a moderately dense environment (p=0.5). (D) When cell wall is highly dense (almost no pores), particle is trapped indefinitely (p=0.8).

**Figure S7.**
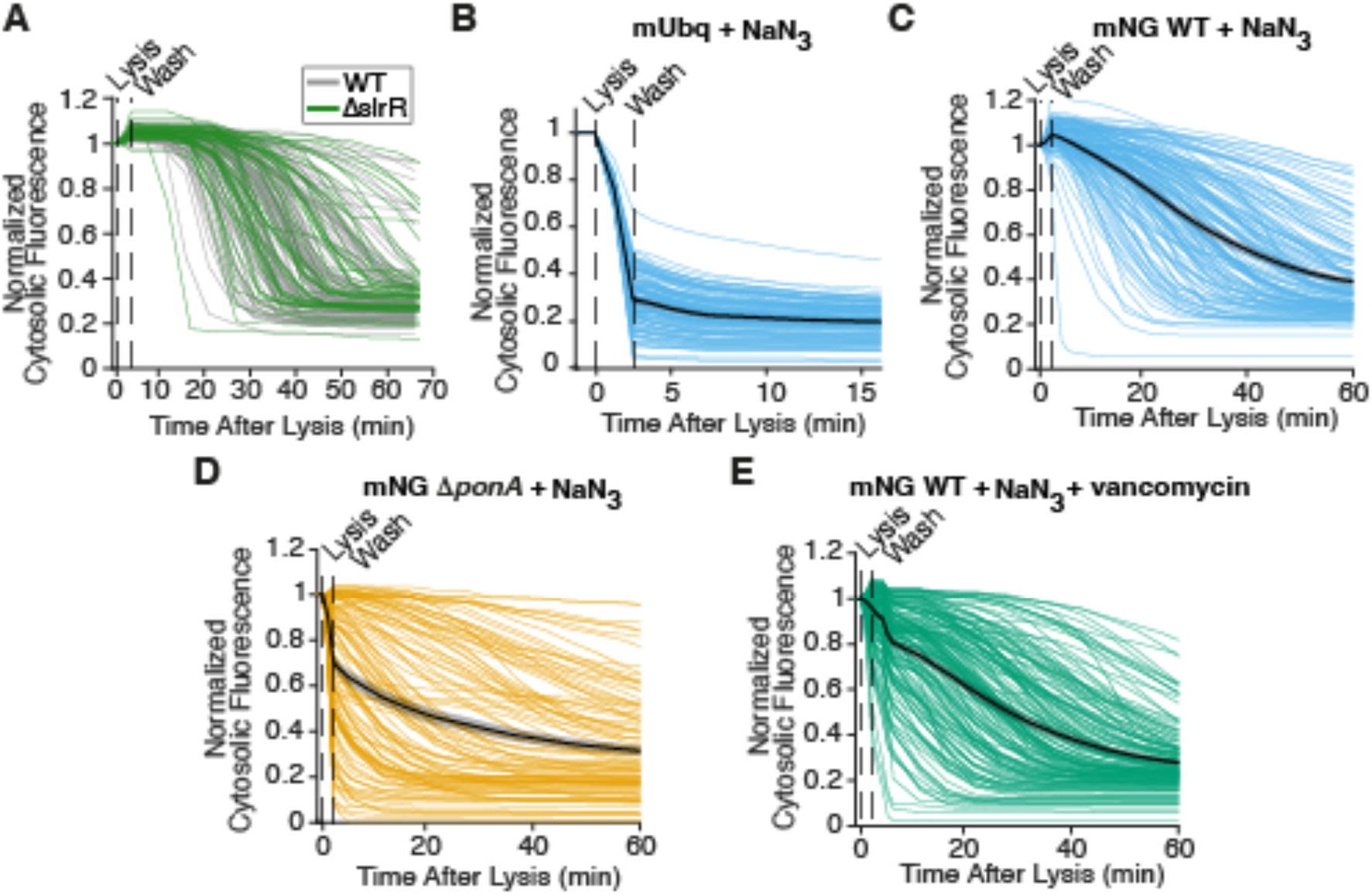
Individual fluorescence traces of sodium azide treated cells. (*A*) Normalized mNeonGreen fluorescence over time from wild-type (WT, gray) and Δ*slrR* (green) cells after detergent treatment in microfluidic perfusion chambers. Shown here for comparison, the untreated data are from the experiments in Fig. 2C. *N* = 1 experiment, *n* = 50 cells. (*B*) Normalized mono-ubiquitin (mUbq) fluorescence traces from the WT + 60 mM sodium azide data shown in Fig. 4F. Mean shown in black. Shaded region indicates standard error of the mean. (*C*) Normalized mNeonGreen (mNG) fluorescence traces from the WT + 60 mM sodium azide data shown in Fig. 4G. Mean shown in black. Shaded region indicates standard error of the mean. (*D*) Normalized fluorescence traces of the Δ*ponA* + 60 mM sodium azide mNeonGreen data shown in Fig. 5D. Mean shown in black. Shaded region indicates standard error of the mean. (*E*) Normalized fluorescence traces of the WT + 60 mM sodium azide + 10 μg/mL vancomycin mNeonGreen data shown in Fig. 5E. Mean shown in black. Shaded region indicates standard error of the mean.

**Figure S8.**
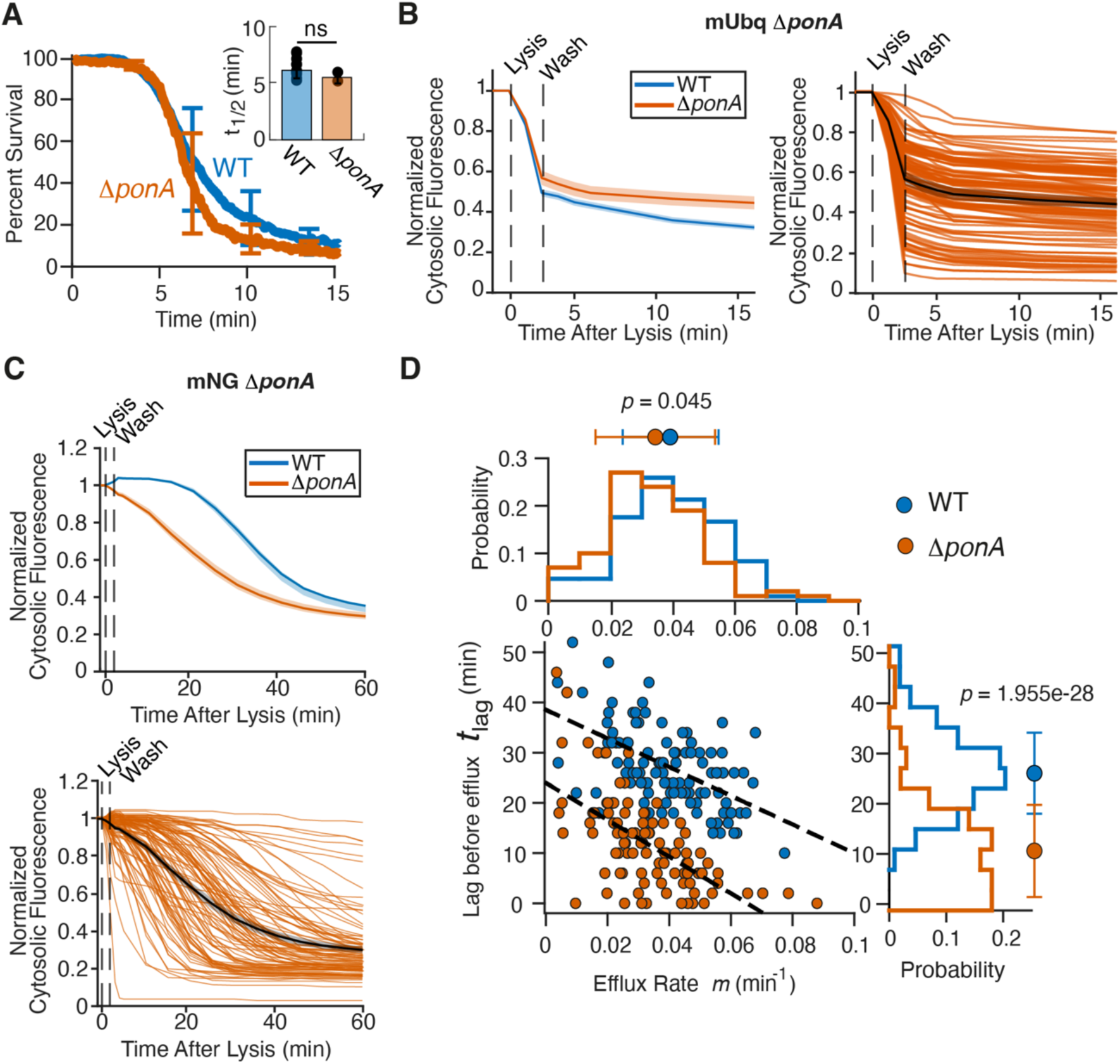
Deletion of PBP1 slightly increases cell wall permeability. (*A*) Mean percent survival over time for Δ*ponA* (red) treated with phospholipase. The wild-type (WT, blue) and Δ*ponA* data are from the experiments in Fig. 1F and Fig. 5A, respectively. Statistical significance calculated using Student’s two-sided *t*-test, ns: p > 0.05. (*B*) Mean (left) and individual (right) normalized mono-ubqiutin (mUbq) fluorescence traces from *ΔponA* (red). Shown here for comparison, the WT mUbq data (blue) from Fig. 2G. Left, shaded region is the standard error of the mean. Right, the black trace is the mean normalized fluorescence. *n* = 115 cells across *N* = 5 technical replicates for *ΔponA* cells. (*C*) Mean (top) and individual (right) normalized fluorescence traces of *ΔponA* mNeonGreen (red) from Fig. 5D. Shown here for comparison, the WT mNeonGreen data (blue) from Fig. 2C. Top, shaded region is the standard error of the mean. Bottom, the black trace is the mean normalized fluorescence. (*D*) Lag vs efflux rate for WT (blue) and Δ*ponA* (red) cells. Histograms of lags (right) and efflux rates (top) show the mean (circles) and standard deviation (error bars).

**Figure S9.**
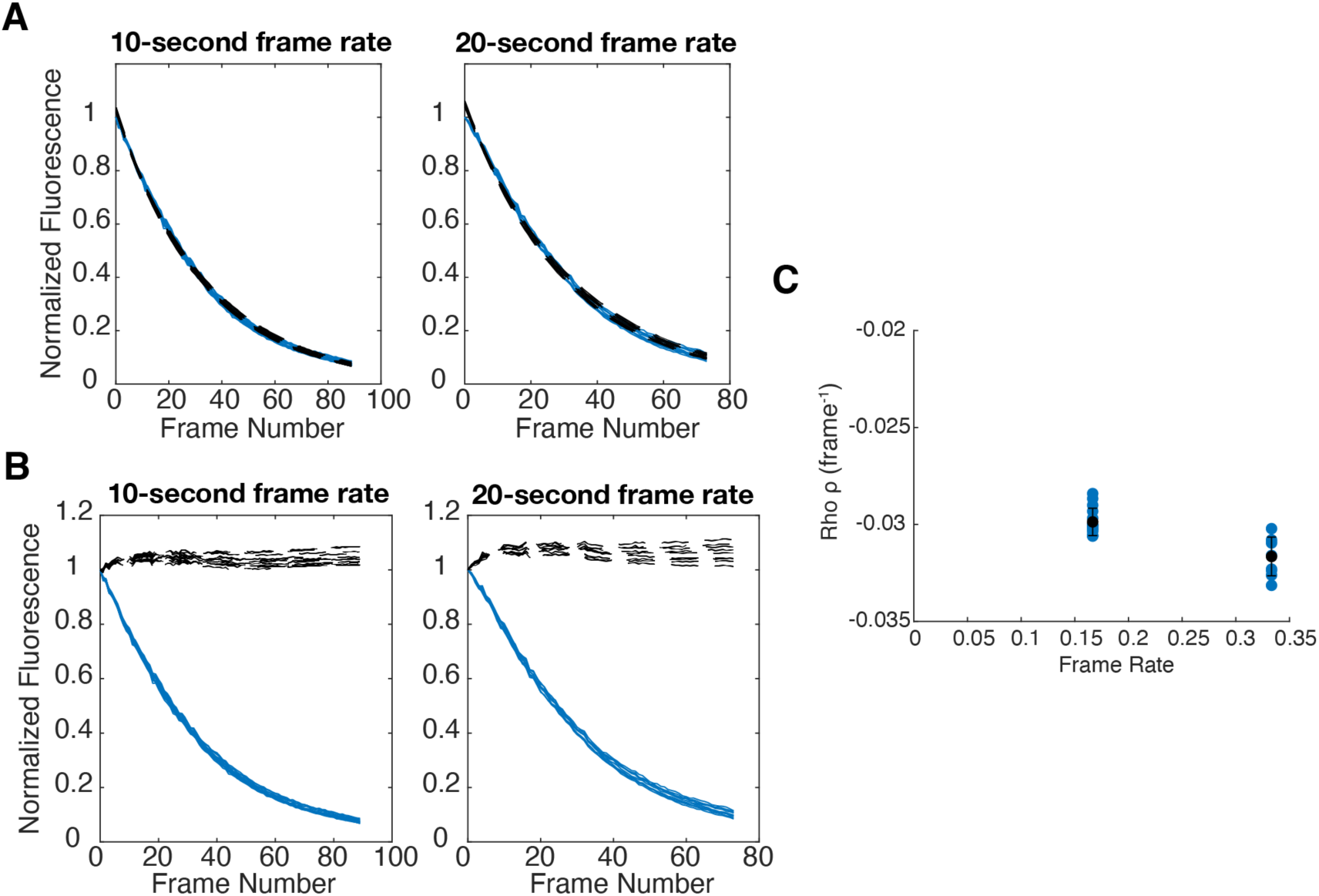
mNeonGreen photobleaching controls. *(A)* The normalized fluorescence traces (blue) fit a single exponential, equation given in the methods (black dashed line). *(B)* The corrected traces (black dashed lines) are obtained by adding the integral of the bleached fluorophore, as given by the photobleaching constant rho (π), to the normalized fluorescence traces (blue). *(C)* Based on the rho vs frame rate, we set the photobleaching constant at 0.03. Each experiment was performed once and *n* = 10, 8 cells for the 10-second and 20-second frame rate experiment, respectively.

**Figure S10.**
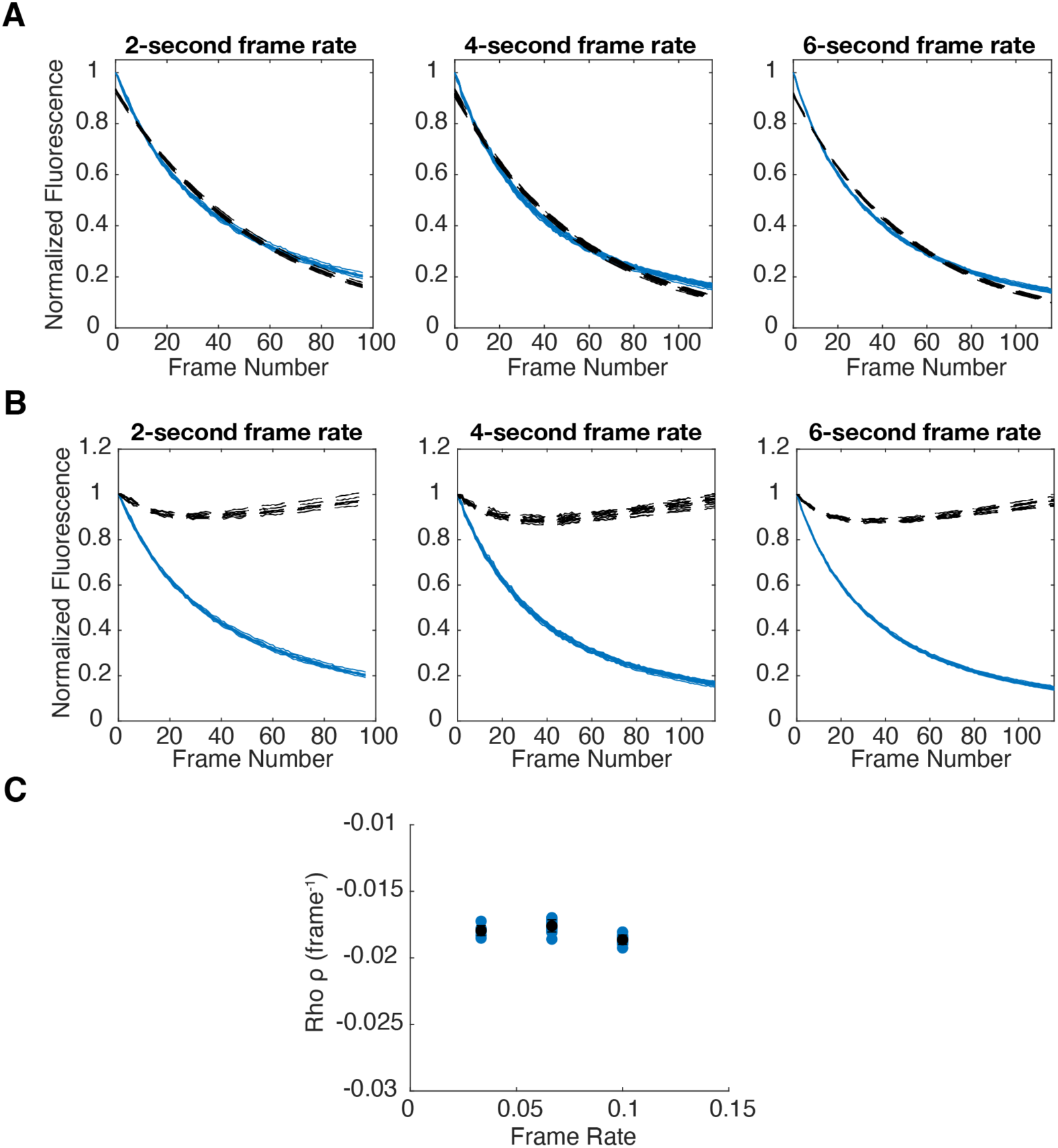
FlAsH-EDT_2_ photobleaching controls. *(A)* The normalized fluorescence traces (blue) fit a single exponential (black dashed line), equation given in the methods. *(B)* The corrected traces (black dashed lines) are obtained by adding the integral of the bleached fluorophore, as given by the photobleaching constant rho (π), to the normalized fluorescence traces (blue). *(C)* Based on the rho vs frame rate, we set the photobleaching constant at 0.02. Each experiment was performed once with *n* = 8, 13, 9 cells for the 2-second, 4-second, and 6-second frame rate experiment, respectively.

**Table S1.**
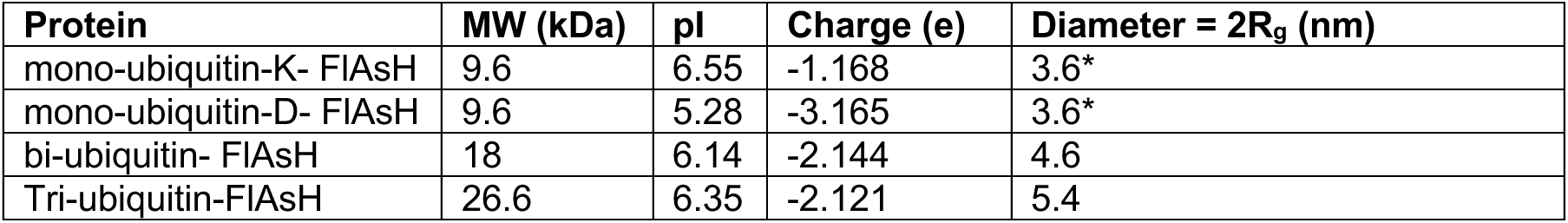
Molecular Properties of Protein Probes (continued). The PDB structures for these proteins were not generated and so the radius of gyration is assumed to be the same as mono-ubiquitin.

| Protein | MW (kDa) | pI | Charge (e) | Diameter = 2R<sub>g</sub> (nm) |
| --- | --- | --- | --- | --- |
| mono-ubiquitin-K- FIAsh | 9.6 | 6.55 | -1.168 | 3.6* |
| mono-ubiquitin-D- FIAsh | 9.6 | 5.28 | -3.165 | 3.6* |
| bi-ubiquitin- FIAsh | 18 | 6.14 | -2.144 | 4.6 |
| Tri-ubiquitin-FIAsh | 26.6 | 6.35 | -2.121 | 5.4 |

**Table S2.** Simulation parameters.

| Parameter | Value | Fig. 3A and Fig. S6 |
| --- | --- | --- |
| Diffusion coefficient, $D$ | 0.81 $\mu\text{m}^2/\text{s}$ | - |
| Lattice constant, $a$ | 3.6 nm | 1.8 nm |
| Time step size, $\Delta t$ | 0.5 $\mu\text{s}$ | - |
| Cell radius, $R$ | 360 nm | 180 nm |
| Wall thickness, $\delta$ | 28.8 nm | 36 nm |
| Cell height, $h$ | 3600 nm | 360nm |
| Step size, $\Delta l$ | 0.9 nm | 0.9 nm |
| Particle diameters, $d$ | 3.6, 4.5 nm | 1.8 nm |
| Wall density, $p$ | 0-1 | 0-1 |
| Simulation length, $N_{\text{steps}}$ | $10^{8.7}$ (in 3B), $10^{9.38}$ (in 3C) | $10^5$ |
| Number of wall realizations | Up to 120 (in 3B), 180 (in 3C for size = 5 nm), and 24 (in 3C for size = 5nm) | - |
| Trajectories per wall realization | 100 | - |

## Notes

### Competing Interest Statement

The authors have declared no competing interest.

